# Brain networks subserving functional core processes of emotions identified with componential modelling

**DOI:** 10.1101/2020.06.10.145201

**Authors:** Gelareh Mohammadi, Dimitri Van De Ville, Patrik Vuilleumier

## Abstract

Emotions have powerful effects on the mind, body, and behavior. Although psychology theories emphasized multi-componential characteristics of emotions, little is known about the nature and neural architecture of such components in the brain. We used a multivariate data-driven approach to decompose a wide range of emotions into functional core processes and identify their neural organization. Twenty participants watched 40 emotional clips and rated 119 emotional moments in terms of 32 component features defined by a previously validated componential model. Results show how different emotions emerge from coordinated activity across a set of brain networks coding for component processes associated with valuation appraisal, hedonic experience, novelty, goal-relevance, approach/avoidance tendencies, and social concerns. Our study goes beyond previous research that focused on either categorical or dimensional emotions and highlighting how novel methodology combined with componential modelling may allow emotion neuroscience to move forward and unveil the functional architecture of human affective experiences.

## Introduction

Emotions are complex and multifaceted phenomena that do not only generate rich subjective feeling states, but also powerfully impact on perception^1^, cognition^2^, memory^3,4^, and action^5^. Various theories have been proposed to characterize emotions and their differentiation, but all remain debated and their links to specific brain processes are still equivocal^6–8^.

Neuroscientific approaches have generally considered emotions either as separate entities (e.g., fear, joy) following theoretical models of discrete emotions^9^ or instead postulated a few basic dimensions (e.g., valence, arousal) following dimensional models^10^. In both cases, the neural substrates of particular emotion categories or dimensions are usually assigned to dedicated brain areas or circuits (e.g., amygdala, striatum)^8,11^. However, these approaches do not easily account for the rich variety of emotions and their anatomical overlap across distributed brain regions as shown repeatedly by neuroimaging studies in the last two decades^11–14^. Several meta-analyses indicate there is no simple one-to-one association of particular emotions or dimensions with individual brain regions^15–17^. Conceptual constructs of valence^18,19^ or arousal^20,21^ do not correspond to clearly separable or unique neural substrates. Therefore, the exact role of different brain areas and their functional interaction within large-scale networks during emotional experience remains unresolved.

In contrast, psychology theories have highlighted the existence of multiple componential processes (e.g. appraisal or action tendency) that interact with each other to evaluate the meaning of events and induce adaptive changes in behavior and cognition^22^. Such models make an explicit distinction between different constituents of emotions, for example, appraisal mechanisms, motor expression, action tendencies, peripheral autonomic changes, motivational drive, as well as various effects on cognitive and memory functions, in addition to the generation of subjective feeling states^23^ (see Figure 1). These constituents may be shared between different emotions but engaged in different ways and to different degrees. Moreover, different appraisal mechanisms evaluate events along different dimensions, which eventually determine their affective significance and trigger corresponding changes in mental and bodily functions^24^. The pattern of appraisals and corresponding responses will, in turn, generate a particular emotion experience. According to such models, appraisals may encompass not only pleasantness (valence), but also novelty, relevance to current goals, causality and agency, expectation and familiarity, control ability, as well as personal values, social norms, and other contextual factors^6^.

**Figure 1:**
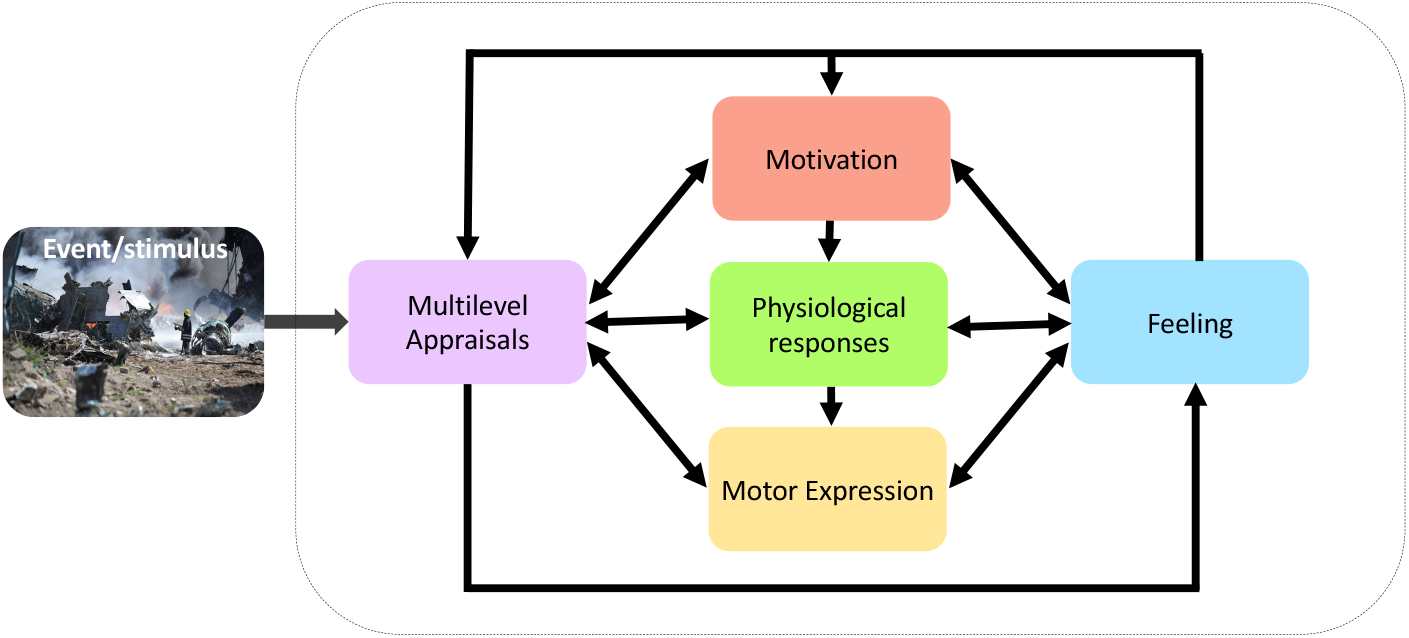
Component model of emotion. In this framework, emotions are conceived as resulting from the concomitant (or sequential) engagement of distinct processes, responsible for the evaluation as well as the behavioral and bodily responses to particular events. According to the Component Process Model (CPM) proposed by Scherer and colleagues, from which emotion features were defined in our study, five distinct functional components are dynamically activating and reciprocally interacting to constitute an emotional experience, including appraisal mechanisms that process contextual information about the event, motivational mechanisms that promote goal-oriented behaviors and cognitions, motor expressions and physiological changes that instantiate bodily responses, and subjective feelings which may reflect an emerging component encoding conscious emotion awareness.

While componential and appraisal theories of emotion have been explored in detail in psychology, no study has examined how these component processes are represented in the brain. However, neuroimaging results show activations in overlapping and distributed regions in response to various emotions^15–17^, consistent with the notion that emotions recruit multiple cognitive or sensorimotor processes subserving their adaptive function. Such activation patterns would appear more consistent with a componential architecture, engaging parallel processes mediated by large-scale brain networks, rather than a modular organization.

Another key issue for emotion studies in psychology and neuroscience concerns the elicitation procedures used to induce emotional responses. Most employ highly simplified and indirect approaches, for example, pictures of facial expressions, voices, music, or photographs that convey only a few specific emotions^8^. These stimuli may cause a perception or recognition of particular emotions, but do not necessarily prompt a genuine emotional experience. Further, when explored with more naturalistic procedures (e.g., movies or memory scripts), emotions are most often studied in terms of pre-defined categories^25^ or dimensions^26^, rather than functional component processes.

Our study addresses these problematic gaps in emotion research by 1) employing a naturalistic and ecologically valid emotion elicitation procedure with a large movie dataset; 2) decomposing emotions into a multidimensional space organized along distinct component processes, based on participant’s experience rather than pre-defined categories; and 3) dissecting the main “building blocks” of this space and their neural substrates using a data-driven modelling approach. Our main goal is to refine our understanding of emotion processing in the human brain through defining new methods to uncover their neural organization. We base our approach on a well-established component process model (CPM) of emotion^22,27^, which provides a comprehensive representation of several key aspects of emotional behavior and experience (see Methods). Our results demonstrate how emotions may emerge from coordinated activity across a distributed set of brain networks coding for component processes associated with valuation appraisal, novelty, hedonic experience, goal-relevance, approach/avoidance tendencies, and social concerns. In doing so, our study goes beyond previous research in several ways and opens new perspectives in affective neuroscience. Figure 2 illustrates the pipeline of our approach (see method section for details).

**Figure 2:**
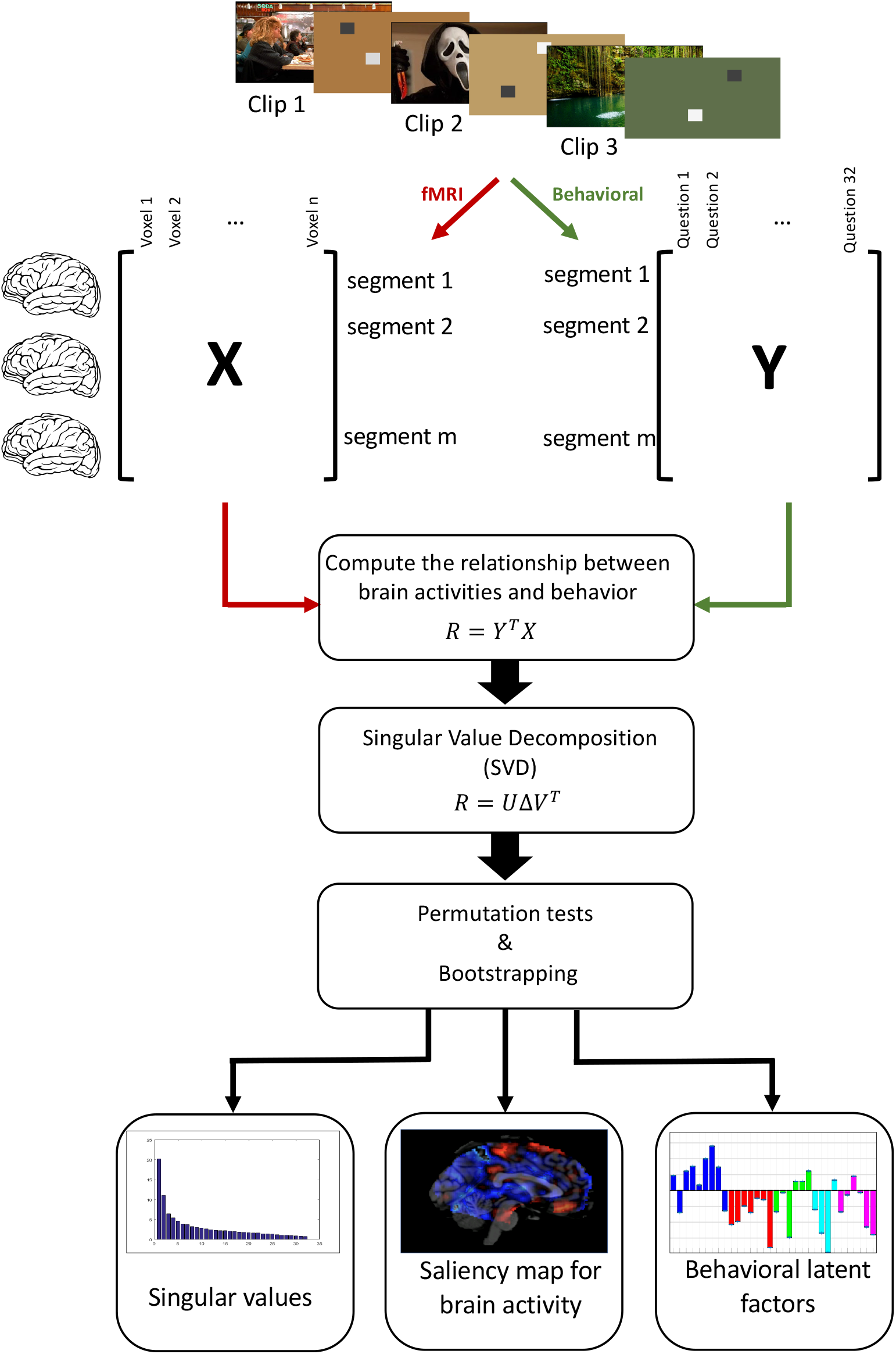
Partial Least Square Correlation (PLSC) method. Participants watch emotional clips during four daily fMRI sessions. Matrix X summarises the brain activity patterns during emotional events of video clips and Matrix Y summarised the assessment of 32 emotion features collected during a separate behavioral session and 2 physiology features for each emotional event. PLSC is then applied to find the commonalities between neural activity and behavioral measures. Th is is achieved in three steps, first by computing the relationship (R) between brain activities (X) and behaviors (Y), followed by decomposing the relationship matrix R using singular vector decomposition (SVD) which allows for projecting both the BOLD fMRI respons e s (X) and behavioral ratings of component features (Y) onto new spaces U and V (latent structures) where the covariance is maximal. Finally, permutation tests and bootstrapping are applied to assess the statistical significance and saliency of each latent factor.

## Results

Our main study presented a group of healthy volunteers (n=20) with a series of 40 video clips, which were selected from a large dataset in a preliminary study (n= 139) so as to convey a large range of emotions (see Methods). Participants watched these movies (mean duration = 111s, SD = 40s) while whole brain activity was measured with fMRI and peripheral physiology was monitored through electrodermal activity (EDA), heart rate (HR), and respiration rate (RR). Movies were presented over 4 different sessions on separate days (10 movies in each, total duration = 74 minutes). During the movies, participants were encouraged to get absorbed in the scenes and let their emotions freely flow without any particular task. After each session, they were presented with the same videos where particular excerpts containing a salient or emotional event (n=1 to 4 excerpts per clip, duration = 12s) were highlighted for assessment following a video pause. Participants had to rate these events in terms of several dimensions of emotional experience (using a 7-point Likert scale). These ratings included 1) a series of 32 descriptors corresponding to major emotion features identified by component models (GRID items, see Table 1) and validated by previous psychology research across several cultures^28^; and 2) a list of 10 discrete emotion categories that could occur during movies (fear, anxiety, anger, sadness, disgust, joy, satisfaction, surprise, love, and calm). We subsequently used the 32 emotion features and two peripheral physiology measures of HR & RR (EDA was also collected, but excluded due to high volume of missing values) to identify consistent covariations corresponding to coordinated patterns of emotional responses, and then applied a multivariate modelling approach to relate each of these patterns to distinctive brain network activations. We also examined how these different components were modulated according to classic emotion categories and dimensions.

**Table 1:** Items of GRID questionnaire. The left column shows the list of the 32 GRID items rated by participants for each of the 119 selected events in movies and two peripheral physiology measures (marked with asterisks). In this column, the abbreviation “s.th.” stands for “something”. The second column represents the component that each GRID item belongs to. The 6 right columns correspond to the six factors obtained from a factorial analysis (D1-D6) and their loading coefficients. Loading coefficients with maximum values for each factor are color-coded to help interpretation and group them in terms of corresponding factors: D1 can be interpreted as valence, D2 corresponds to arousal, D3 mainly encodes motor expression and bodily changes, D4 represents novelty, D5 relates to action tendencies, and finally D6 represents norms. Coefficients highlighted in bold are statistically significant (p-value <.05).

| Questions: (While watching this scene, did you...) | Components | D1 | D2 | D3 | D4 | D5 | D6 |
| --- | --- | --- | --- | --- | --- | --- | --- |
| feel good? | Feelings | **-0.85** | -0.12 | -0.17 | -0.06 | -0.13 | -0.03 |
| feel situation was unpleasant for you? | Appraisal | **0.81** | 0.20 | 0.29 | 0.01 | 0.10 | 0.11 |
| feel bad? | Feelings | **0.81** | 0.27 | 0.25 | 0.01 | 0.09 | 0.06 |
| want to undo what was happening? | Motivation | **0.81** | 0.21 | 0.15 | 0.01 | 0.15 | 0.17 |
| want the situation to continue? | Motivation | **-0.81** | 0.12 | -0.05 | -0.07 | -0.11 | -0.02 |
| feel the urge to stop what was happening? | Motivation | **0.81** | 0.19 | 0.17 | 0.00 | 0.16 | 0.16 |
| feel it was unpleasant for someone else? | Appraisal | **0.71** | 0.13 | 0.07 | 0.18 | 0.16 | 0.18 |
| feel calm? | Feelings | **-0.69** | -0.19 | -0.26 | -0.11 | -0.18 | -0.08 |
| feel strong? | Feelings | **-0.54** | 0.04 | -0.08 | -0.08 | -0.01 | 0.03 |
| want to tackle the situation and do s.th.? | Motivation | **0.51** | 0.35 | 0.00 | -0.05 | 0.38 | 0.14 |
| feel an intense emotional state? | Feelings | 0.19 | **0.82** | 0.28 | -0.07 | 0.06 | 0.06 |
| experience an emotional state for a long time? | Feelings | 0.17 | **0.79** | 0.24 | -0.06 | 0.06 | 0.04 |
| feel motivated to pay attention to the scene? | Motivation | -0.15 | **0.42** | -0.08 | 0.06 | 0.08 | 0.05 |
| have a feeling of lump in the throat? | Physiology | 0.42 | **0.40** | 0.29 | -0.09 | 0.14 | -0.02 |
| show tears? | Expression | 0.15 | **0.39** | 0.01 | -0.12 | 0.08 | -0.13 |
| think it was important for somebody's goal | Appraisal | 0.06 | 0.18 | 0.06 | 0.01 | 0.00 | 0.03 |
| experience muscles tensing? | Physiology | 0.45 | 0.21 | **0.49** | 0.01 | 0.22 | 0.04 |
| produce abrupt body movement? | Physiology | 0.19 | 0.10 | **0.48** | 0.10 | 0.19 | 0.09 |
| close your eyes? | Expression | 0.23 | 0.05 | **0.45** | 0.00 | 0.05 | 0.01 |
| have stomach trouble? | Expression | 0.35 | 0.37 | **0.45** | -0.01 | 0.10 | -0.04 |
| feel warm? | Physiology | 0.02 | 0.15 | **0.39** | -0.02 | 0.10 | -0.05 |
| have eyebrow go up? | Expression | 0.15 | 0.04 | 0.30 | 0.27 | 0.07 | 0.20 |
| press lips together? | Expression | 0.26 | 0.24 | 0.28 | 0.01 | 0.05 | 0.05 |
| have the jaw drop? | Expression | 0.06 | 0.09 | 0.23 | 0.26 | 0.10 | 0.12 |
| *Heart rate | Physiology | -0.01 | 0.02 | 0.20 | 0.03 | -0.06 | 0.02 |
| *Respiratory rate | Physiology | 0.09 | -0.04 | 0.15 | 0.05 | 0.05 | 0.06 |
| feel that the event was unpredictable? | Appraisal | 0.08 | -0.02 | 0.09 | **0.77** | 0.01 | 0.06 |
| feel the event occurred suddenly? | Appraisal | 0.21 | 0.03 | 0.13 | **0.68** | 0.05 | 0.06 |
| think that the consequence was predictable? | Appraisal | 0.06 | 0.07 | 0.02 | **-0.42** | 0.09 | 0.05 |
| think the event was caused by chance? | Appraisal | 0.01 | -0.04 | -0.01 | **0.42** | -0.04 | -0.01 |
| want to destroy s.th.? | Motivation | 0.30 | 0.15 | 0.20 | -0.06 | **0.73** | 0.05 |
| want to damage, hit or say s.th. that hurts? | Motivation | 0.35 | 0.18 | 0.18 | -0.06 | **0.70** | 0.12 |
| think it violated laws/social norms? | Appraisal | 0.59 | 0.07 | 0.16 | 0.09 | 0.18 | **0.64** |
| think it was incongruent with your standards? | Appraisal | 0.61 | 0.06 | 0.20 | 0.06 | 0.16 | **0.62** |

### Behavioral and physiological measures

To illustrate the variability of elicited emotions, Figure 3a presents the histogram of the discrete emotion categories, selected by our participants as the most dominant emotion during each of the salient movie excerpts. Although the distribution of discrete emotions is not uniform, it shows that, except for love, our stimulus material and experimental design was successful in eliciting a wide range of different emotions, allowing us to obtain a comprehensive survey of the componential space. The non-uniform distribution of emotion categories (unlike results from the movie selection phase; see Methods) is due to the fact that these ratings concerned short emotional episodes (single events) and were obtained from a varying number of segments across different video clips (unlike the more global judgments made for whole movies during pre-selection). Moreover, single events in a movie did not necessarily elicit the same emotion as the global judgment made for an entire movie clip, which highlights the importance of using and characterizing short segments for fMRI analysis.

**Figure 3:**
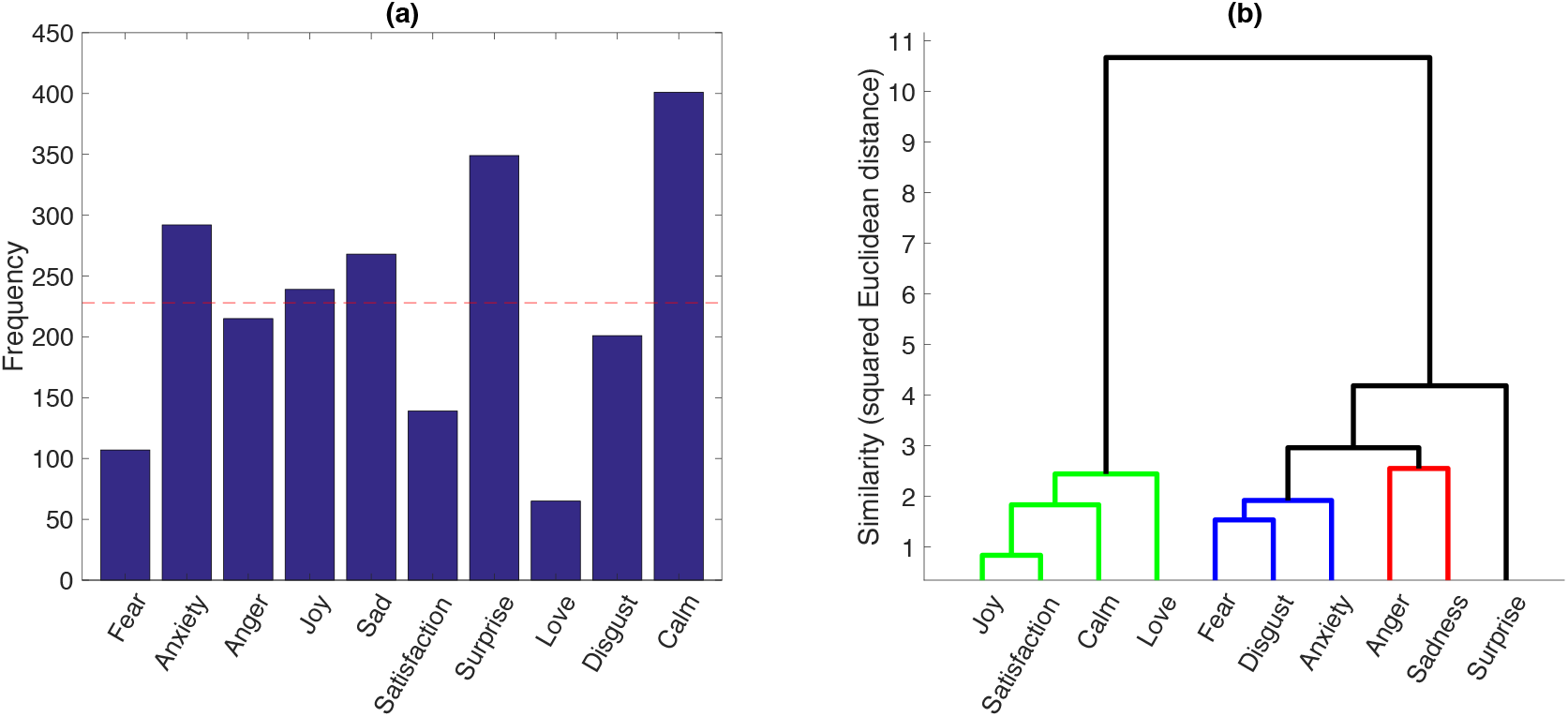
Histogram and hierarchical clustering of discrete emotions. **(a)** Histogram of categorical emotions based on their frequency in the ratings of 119 emotional event by 20 participants (data from 2 participants were not complete). The red dashed line shows the ideal frequency if samples were distributed uniformly. **(b)** Hierarchical clustering of the discrete emotion profiles in the GRID space using Ward algorithm. The higher-level clusters distinguish between positive and negative emotions. The lower-level clusters reflect a segregation of feelings in terms of pleasantness (green), surprise (black), distress (blue), and annoyance or frustration (red).

In addition, a histogram count of the GRID features across movie emotional events (Figure 4) showed that ratings of all 32 items were generally well spread across the two ends of the continuum, with only a few items exhibiting a distribution skewed towards the lower end (rated as 1). This bias was more evident for features from the expression component, most likely due to the passive condition of emotional experience during movie watching that does not require any direct behavioral responses or communication.

Altogether, these behavioral data demonstrate that our procedure could successfully cover the whole componential space for a range of different emotions and thus effectively test for patterns of shared variability across the different stimulus conditions.

**Figure 4:**
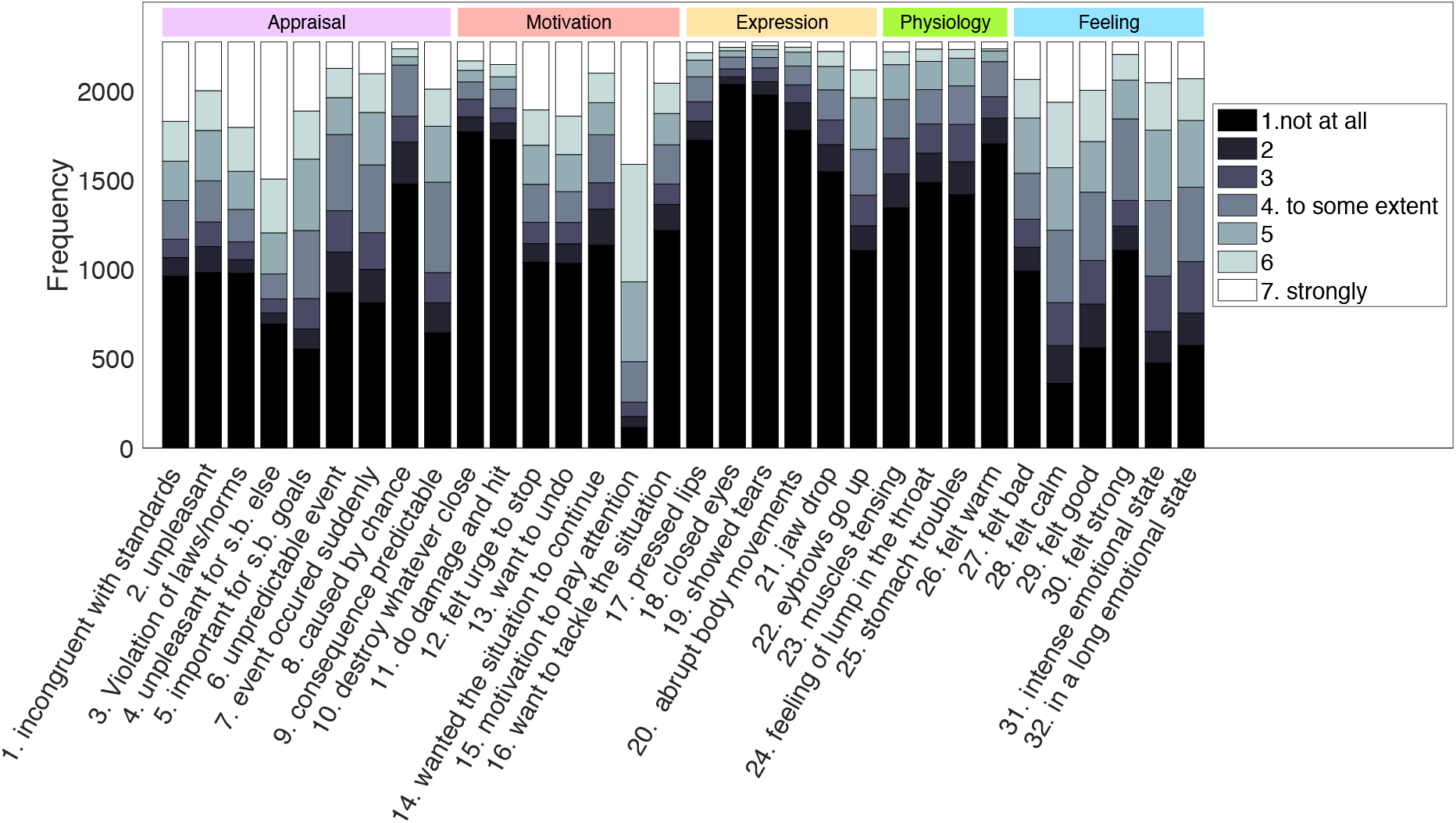
Histogram of GRID items. Histogram of ratings for all the 32 GRID items based on the number of times a specific rating (1-7) was selected across all assessments (119 assessments per participant) and all 20 participants (the data from 2 participants were not complete). The abbreviation “s.b”. stands for somebody.

Finally, physiology recordings obtained during movies (see methods) confirmed significant bodily changes across different emotions for HR (*F*(9,2291) = 3.767, *p* < .001, *η*^2^ = .014), RR (*F*(9,2291) = 3.767, *p* < .001, *η*^2^ = .014), and tonic EDA (*F*(9,2029) = 3.415, *p* < .001, *η*^2^ = 0.014). No reliable effect was found for phasic EDA (*F*(9,2029) = 1.237, *p* = .268, *η*^2^ = .005). Please note we report only basic measures of these physiological signals (mean and variance) as part of GRID descriptors, rather than more sophisticated characteristics used in some behavioral studies^12^, but these were not the main variables of interest in our study and reliable physiological data are difficult to record during concomitant fMRI. However, in a separate analysis (not reported here), we found that the absolute mean and variance of HR, RR, and EDA could predict appraisal GRID descriptors significantly above chance level, while they did not reliably predict discrete emotion ratings. These findings suggest that our measures were robust enough to be used in subsequent fMRI analysis, even though clearly insufficient to fully differentiate complex emotional experiences by themselves (see methods, and supplementary methods, section A). To fully characterise emotional responses in terms of physiology processes, more analyses has to be conducted, which is out of the scope of this study.

### Cluster analysis

To analyse the relation among different emotion categories within the componential model space, we performed a hierarchical clustering analysis on the average profile of each discrete emotion along 32 GRID features plus two peripheral physiology measures of HR & RR. As can be seen in Figure 5, different features were present to different degrees for different discrete emotions, while several features were shared by more than one particular emotion. Accordingly, the clustering results indicated a clear distinction between positive versus negative emotions at the higher level of differentiation (Figure 3b), while four different clusters were observed at the lower level representing, respectively, pleasant feelings (joy, satisfaction, love, calm), distress (fear, disgust, anxiety), annoyance (anger and sadness), and surprise. Surprise showed a higher similarity to negative categories than to pleasantness, possibly reflecting that it came mostly with negative contexts in our movie dataset.

**Figure 5:**
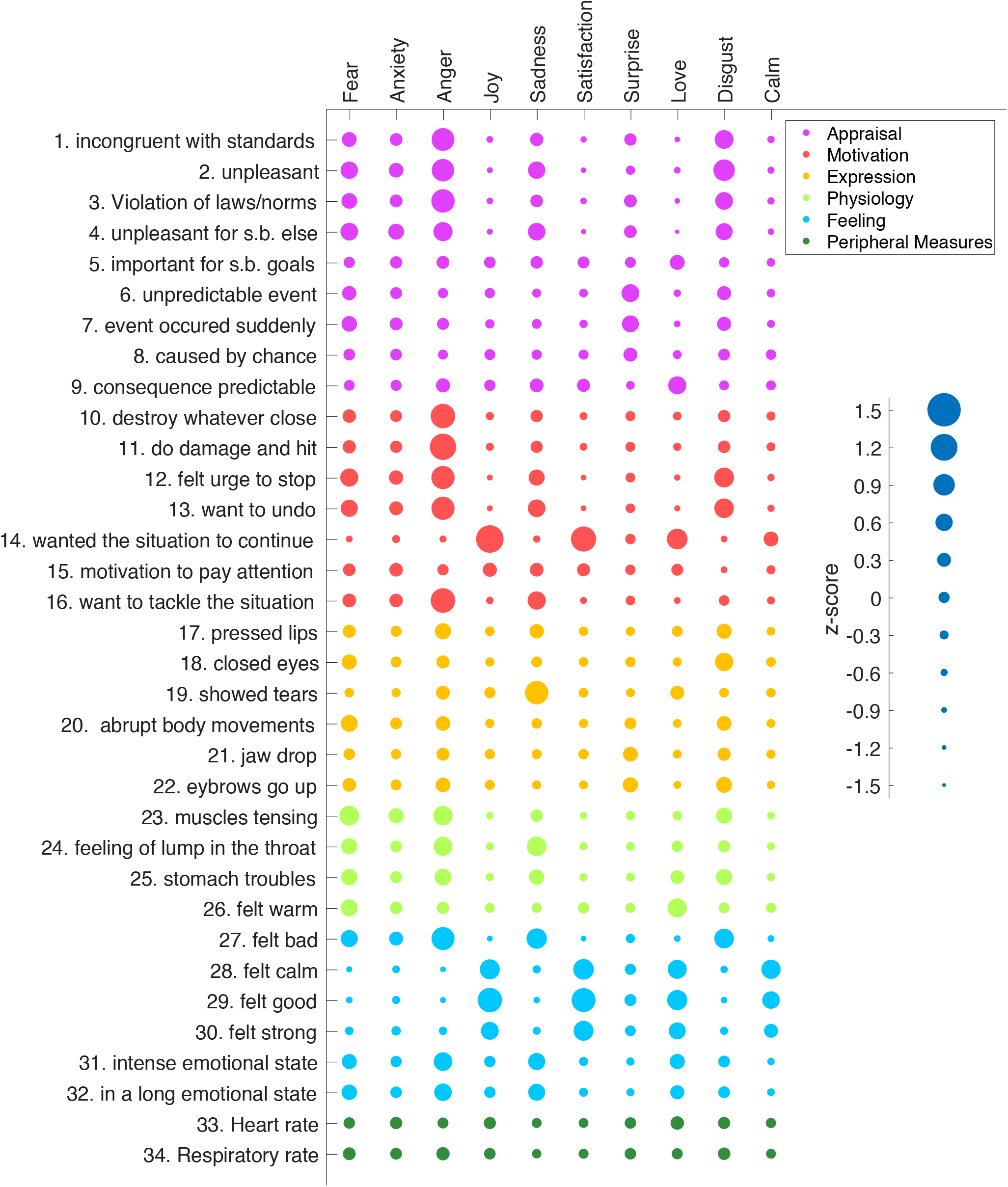
Discrete emotion profiles in GRID space. Average profile of each discrete emotion on the 32 GRID features and two peripheral physiology measures (HR, RR) after within-subject normalisation. For each discrete emotion, all assessments from all 20 participants with that discrete emotion label was used. Each bubble corresponds to a z-score using an exponential scaling. The smallest bubble represents a z-score=-1.27, corresponding to “not at all”, and the biggest bubble represents a z-score=+1.23 corresponding to “felt strongly”. Colors represent the different emotion components to which GRID items belong to.

Altogether, these findings demonstrate that theoretically meaningful clusters can be derived from our set of GRID features, fully in accordance with emotional dimensions commonly considered in neuroscience studies. This in turn further validates our experimental paradigm by showing it could successfully elicit different emotion categories with expected characteristics.

### Underlying factors

For completeness and comparison with prior psychology research^29^, we also performed an exploratory factor analysis to recover common dimensions underlying the different GRID features^28^. Table 1 summarizes these six factors and their relative loadings. Overall, about 47.8% of the total variance was explained by these six factors. The GRID items most strongly associated with these 6 factors suggest they may be linked to pleasantness (21.6%), arousal (7.6%), expression (5.9%), novelty (5.0%), action tendency (4.6%), and social norms (3.1%), respectively. This accords with previous findings obtained with similar component models^29^. These results also confirm that emotional experiences involve more than two dimensions of valence and arousal when explored across a variety of conditions with ecological validity.

Taken together, these behavioral data converge to indicate that a comprehensive differentiation of emotions would require not only to go beyond valence and arousal dimensions, but also to better characterize the commonality and specificity of different types of emotions. Accordingly, this also implies that mapping the neural underpinnings of emotion requires a multivariate approach that transcends two-dimensional representations or simple oppositions between discrete categories, as we propose with the componential approach in the next section.

### Decomposing emotions into functional core processes and their neural substrates

To determine how component processes of emotion are organized in relation to functional brain systems, we collected fMRI data from our 20 healthy participants while they watched the 40 emotional video clips, and then analysed blood-oxygenation-level-dependent (BOLD) time-courses to identify patterns of brain activity corresponding to systematic covariation in their ratings of specific componential features during the movies (see Figure 2). For each participant, we used ratings of each of the 32 GRID features for each of the 119 salient emotional events selected from these movies (total of 2276 surveys, including 72’832 rating scores), plus the HR and RR z-score values for these events. To decompose emotion responses into core functional processes, we applied a multivariate technique, namely Partial Least Square Correlation (PLSC)^30^, enabling us to analyse covariance in two feature spaces: (1) the multidimensional structure of emotion ratings along all GRID items (component process model), and (2) the multidimensional activation patterns across all brain voxels. PLSC identifies a set of orthogonal latent variables for each space that express maximum cross-covariance and thus represent shared information in the two spaces (i.e., behavior and brain). In other words, this method allows for modelling the functional relationship between the coordinated mobilization of multiple emotion features and corresponding variations in neural activity.

After pre-processing fMRI data according to standard pipelines and normalizing behavioral ratings to within-subject z-scores, the PLSC analysis was applied to the whole sample from our 20 participants. Significance of components (p-values) were calculated using permutation tests, and z-scores reflecting reliability of loadings (a.k.a., saliencies) were estimated using bootstrap ratio. To determine the generalisability of the method and overcome sample size limitations, a conservative bootstrap strategy was taken in which, at every bootstrap iteration, only 14 randomly selected subjects were used, and the final behavioral loadings at group-level were estimated as the average loadings across all bootstrap iterations. The standard deviation of each loading was also considered as a measure of stability of the loadings across different sets of subjects and hence provided an estimate for the robustness/generalisability of our results (see methods for details).

The PLSC analysis revealed 6 significant latent variables (LV) with p< .01. These latent variables represent distinct combinations of behavioral features with concomitant brain patterns. Importantly, please note that LVs are interpreted in terms of both their component features and associated neural activity. Figure 6 shows the loading profile along all GRID and physiology features for each latent variable identified here: LV1 (p<.001, 19.0%), LV2 (p<.001, 11.0%), LV3 (p<.001, 6.0%), LV4 (p<.001, 5.1%), LV5 (p<.01, 4.6%), and LV6 (p<.01, 3.4%); numbers in parenthesis indicate p-value and percentage of covariance explained, respectively. Positive or negative loadings of particular features reflect the relative presence or absence of these features, respectively. Their weighted sum represents a specific combination linked to a particular brain activation pattern, whose expression characterizes a given functional core process underlying the generation of an emotion response. All maps were thresholded at positive or negative saliency values of >2.5 or <-2.5 (p<.01). The dominant behavioral profile of LV1 shows higher weightings for several features related to the appraisal of values with motivational aspects and the valence dimension of the feeling component. As illustrated in *Figure 6*, positive ratings for this LV reflect events with low/no unpleasantness, low/no incongruence with standards, a desire to continue/not to stop, and feeling good/not feeling bad (among others). On the brain side, the voxel-wise saliency map of LV1 exhibits significant positive weights (>2.5) in ventromedial prefrontal cortex (vmPFC), lateral orbitofrontal cortex (OFC), central/lateral amygdala and ventral tegmental area (VTA), and significant negative weights (<-2.5) in anterior and dorsal insula, mid cingulate cortex, thalamus, putamen, dorsal amygdala and substantia inominata, as well as medial parietal and lateral occipital areas (Figure 7).

**Figure 6:**
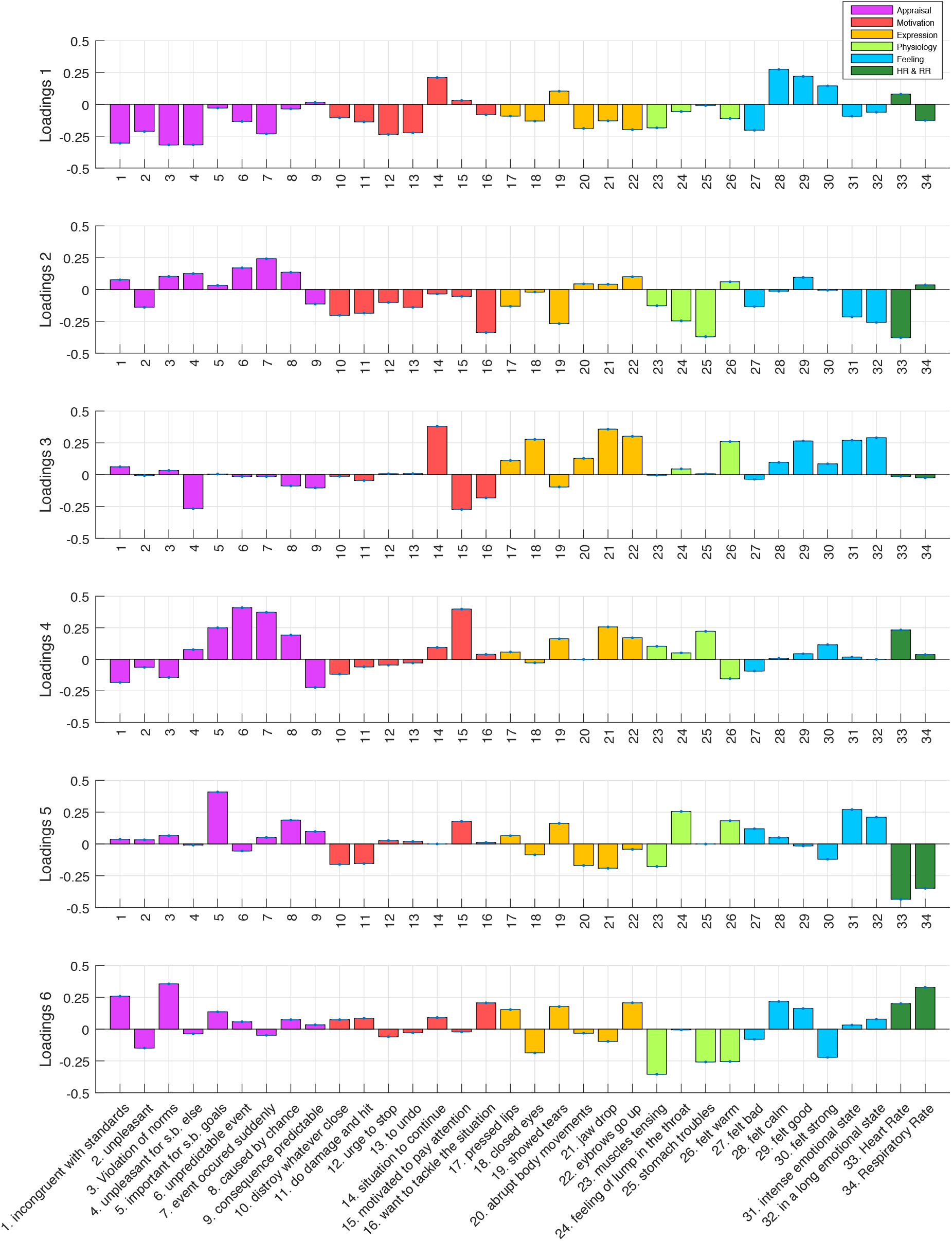
Loadings of partial least square correlation. Loadings of PLSC for GRID items (behavioral) and peripheral measures corresponding to the six significant latent variables (1-6) respectively interpreted as: valence, novelty, hedonic impact, goal monitoring, goal relevance, and avoidance. Each loading vector corresponds to one brain activity map that is shown in Figure 7. The blue dots on each bar indicate the standard deviation that reflect reliability of the loading when apply bootstrapping; however, due to very small variation, they are not very visible.

**Figure 7:**
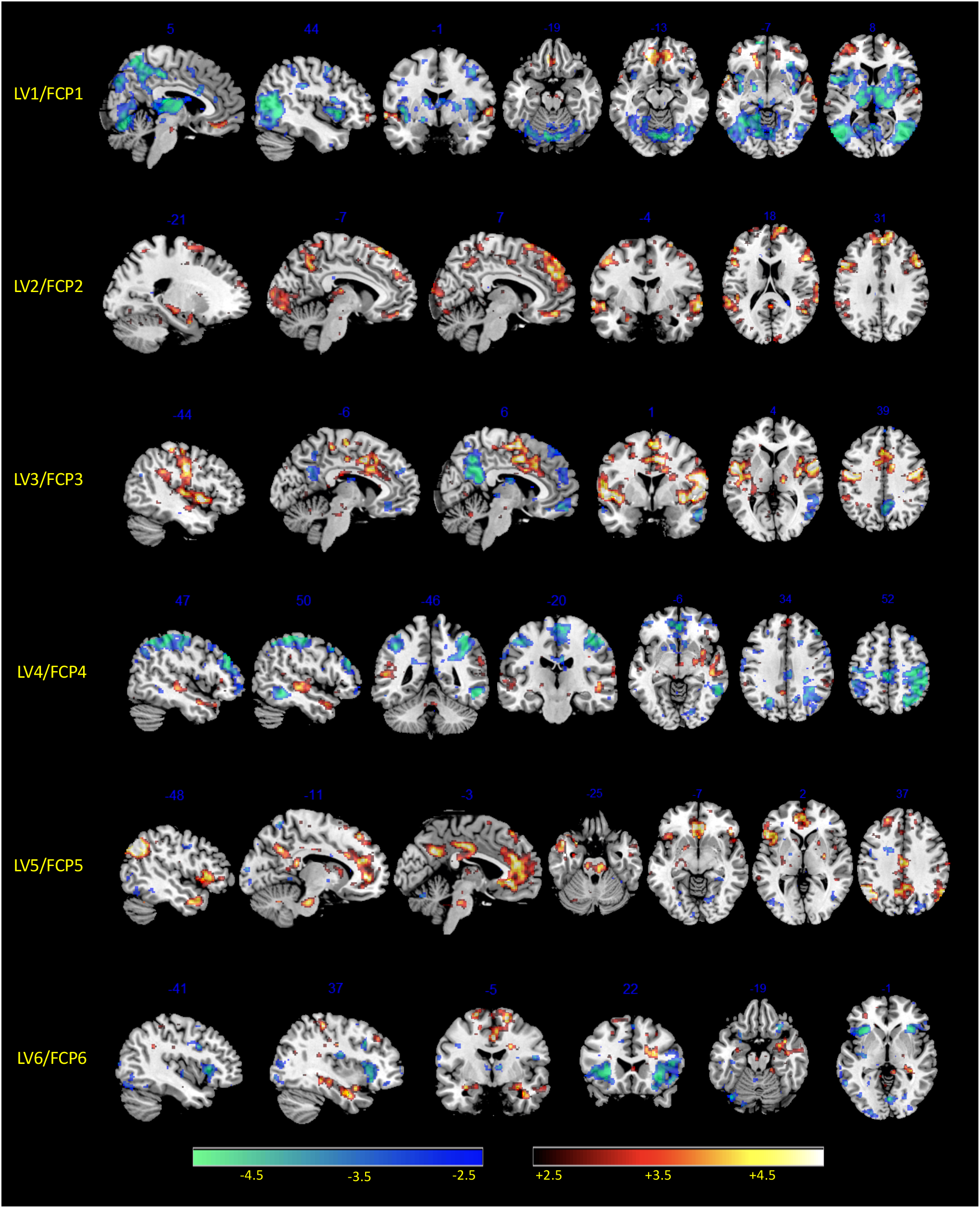
Brain saliency maps. Brain activity maps of relative saliencies corresponding to each of the six significant functional core processes, a.k.a. latent variables (LV), obtained by the PLS analysis of GRID ratings. The red spectrum accounts for positive saliencies above +2.5 and blue spectrum corresponds to negative saliencies below −2.5

The second latent variable LV2 shows higher weightings on items related to appraisals of unexpected event and detection of changes (i.e., sudden and unpredictable, with brief emotion intensity), a pattern that can be interpreted as novelty detection (Figure 6). The corresponding brain saliency maps show significant positive weights bilaterally in dorsomedial prefrontal cortex (DMPFC) and dorsolateral prefrontal cortex (DLPFC), inferior frontal gyrus (IFG), posterior cingulate cortex (PCC), together with large effects in sensory (auditory and visual) cortices, as well as smaller activation clusters in dorsal amygdala, hippocampus, and parahippocampal gyrus (PHG). Negative saliency weights are much weaker and limited, involving only very small parts of anterior insula and rostral ACC (Figure 7).

The third latent variable LV3 loads mainly on expression-related features (such as closing the eyes, pressing the lips, raising eyebrows) along with feeling features related to pleasantness and arousal (warm, good, intense and lasting experience), which together might encode generally pleasurable sensation and hedonic impact. This LV exhibits significant positive saliencies in widespread sensorimotor areas, including the primary somatosensory and motor cortices, particularly over the central fronto-parietal operculum (face area), but also SMA, preSMA, dorsal ACC, posterior insula, ventral tegmental area (VTA), and brainstem (central pons). Negative saliencies were again weak, essentially limited to the PCC, inferior parietal lobule (IPL), and a small sector of ventromedial prefrontal cortex (VMPFC), which may constitute parts of the Default Mode Network (DMN)^31^.

The latent variable LV4 unfolds mainly on appraisal components related to expectation and goal settings, as well as motivated attention and congruence with norms, without any consistent loadings related to feelings. These features may reflect encoding of goals and intentions (of someone else in movies) or violations of expectations. At the brain level, LV4 is associated with positive saliency weights predominating in STS and temporal pole, as well as in DMPFC and precuneus, partly overlapping with social cognition and theory-of-mind networks^32^. There were also smaller clusters in the temporoparietal junction (TPJ), inferior frontal gyrus (IFG), and pallidum, all parts of the ventral attention orienting network^33^. Widespread negative saliency weights were found in superior parietal cortex (intraparietal sulcus, IPS), dorsolateral prefrontal cortex (DLPFC), anterior cingulate cortex (ACC), and posterior SMA, overlapping with dorsal attention networks.

The next significant latent variable LV5 loads selectively on appraisal of event relevance for someone else with some features of physiology (warmth and lump in the throat) and feeling components (high intensity but no valence). There is also a relative lack/suppression of aggressive motivational tendencies (wanting to attack or damage), which, altogether, may reflect a state of prosocial concern or caring for others. The brain saliency map reveals a strong positive correlation with areas of the default mode network (DMN), particularly MPFC and PCC, but also smaller clusters in bilateral anterior insula, inferior parietal lobe, caudate, and brainstem (possibly overlapping with locus coereleus). There are no consistent negative weights.

Finally, the last significant latent variable LV6 loads on anti-social features (e.g., incongruence with norms), attention and approach/attack, but also some degree of pleasant feelings (good or calm), which might relate to curiosity/interest and elements of aggression or active defence. The corresponding brain saliency map shows positive weights in the amygdala, VTA, medial OFC/VMPFC and superior frontal gyrus (SFG), as well as somatosensory areas and PCC (Figure 7); whereas negative saliencies were mainly found in the bilateral anterior insula, lateral OFC, rostral ACC, and posterior visual areas (Figure 7). A summary of average saliency values across cortical areas for each LV is provided in supplementary results, section A, Supplementary Table 3.

The 6 LVs also disclosed distinct patterns of loadings for peripheral physiology measures (Figure 6, last two columns, dark green). Both HR and RR exhibited negative weights for LVs 5 and 6, whereas LV2 was associated with negative weights and LV4 with positive weights of HR alone (not RR). Both measures were only weakly modulated by LVs 1 and 3. These physiology patterns seem compatible with psychological processes putatively associated with each LV^34^. However, similar LVs were obtained when repeating our PLSC analysis without physiological measures, suggesting the latter did not make a major contribution to the results.

Taken together, the joint interpretation of behavioral loadings and their associated brain activation patterns points to distinct functional core processes (FCP) that were mobilized in a coordinated manner and constitute major “building blocks” of the various emotions elicited by the movies.

In order to further investigate the functional role of latent variables identified above (Figure 6) and examine their similarity with discrete emotion categories, as rated by participants for the same event (as in Figure 5), we performed a correlation analysis (Pearson coefficient) to relate each discrete emotion profile to the 6 different LVs. This analysis highlighted that each of the different LVs identified by our data-driven PLSC analysis contributed to different emotions, but to variable degrees, and also they generally held meaningful relationships with discrete categorical labels (see supplementary results, section B).

## Discussion

Our study demonstrates a new data-driven approach to uncover the neural substrates of emotion without imposing any a priori or predefined categories, dimensions, or stimulus classes. Instead, our experimental design relies on solid theoretical models and established concepts of emotion in psychology which go beyond dimensional or categorical models. Our results reveal at least six distinct functional core processes (FCP), engaged in response to emotional experiences during naturalistic movies, and denoted as latent variables (LV 1-6) linking particular combinations of emotion features with particular patterns of brain-wide activity. The six FCPs identified here appear to encode appraisal of value, novelty, hedonic experience, goal monitoring, caring for others, and approach tendency/curiosity. Their neuroanatomical underpinnings involve distributed brain networks, including regions consistently implicated in emotion processing (e.g., amygdala, VMPFC, insula, VTA) and also other regions (e.g., associated with sensorimotor function, social cognition, attentional control), which accord well with appraisal theories and componential models of emotion^22,28^. Importantly, each FCP contributes to different emotion types independently and to different degrees; hence, one particular FCP can be mobilized in more than one emotion as illustrated by strong correlations observed between individual FCPs and specific discrete emotions. We also found that some of the FCPs have higher loadings on one particular component (e.g., appraisal or feeling) but most often include mixed features that do not purely correspond to one unique component as traditionally distinguished in CPM (see Figure 1). This may reflect a limitation of subjective ratings that fail to capture pure componential processes, or more fundamental principles of brain function whereby any cognitive or affective task emerge from the interaction of multiple neural systems.

The functional core processes identified with the current methodology bear several similarities with basic dimensions considered in past research (such as valence or arousal)^10,29^, but also differ in important ways that offer novel insights into the componential nature of emotions. FCP-1 clearly resembles a bipolar valence dimension, often highlighted as the dominant aspect of emotional experience across many studies^10^. Here FCP-1 related bipolar elements of motivational values and feelings with brain regions implicated in affective evaluative processes (such as amygdala and VTA) and affect-based decision making (such as VMPFC, OFC and insula). Mobilization of this network is tightly associated with positive versus negative emotion categories, and thus consistent with valence being a major constituent of core affect^35^. However, an involvement of action features (e.g., urge to stop) and brain areas (e.g., VMPFC) associated with motivation and decision making, but low loading on feeling intensity, suggest that FCP-1 may relate to the theoretical construct of “wanting” (or not wanting)^36^, rather than to the hedonic experience of “liking”^36^.

In contrast, FCP-3 is also characterized by elements of pleasantness, but with high intensity feeling features and motor expression features, suggesting a more direct implication in hedonic experience, reminiscent of the “liking” rather than “wanting” aspect of valence^36^. Accordingly, this FCP shows strong positive correlation with joy and love, but negative correlation with sadness and anxiety, like FCP-1^36^. Neural activation in regions overlapping with motor face areas in operculum and preSMA would be consistent with networks controlling smiling and laughing^37^. Electrical stimulation of medial premotor cortices may elicit both laughter and mirth^37^, and covert activation of zygomatic muscles is a reliable marker of positive valence^38^. Unfortunately, due to technical limitations, facial EMG was not collected in the MRI scanner. Our findings of two distinct networks (FCP-1 and FCP-3) correlating with valence features converge with growing evidence that this dimension may not have a unique psychological or neural underpinning^19,39^.

Another core process, FCP-2, appears to selectively encode novelty, with several memory-related areas, sensory areas, and prefrontal areas. The amygdala is also engaged, in keeping with its role in novelty detection^40^. This FCP is highly correlated with ratings of surprise and may constitute a key substrate for this non-valenced emotion. A prominence of novelty neatly accords with theories of emotion that place it as a frequent dimension after valence and arousal^29^, or an essential appraisal initiating emotion episodes^22^. Our data suggest that novelty detection is not only an important constituent of emotion but also critically relies on memory functions evaluating contextual information. Notably, however, FCP-2 is low on action features and feeling intensity, suggesting it does not reflect a dimension of arousal or alertness.

The FCP-4 bore similarities with FCP-2 but with higher loadings on appraisals of goals, expectations and attention, which covaries with activation of brain networks mediating social cognition and attentional reorienting mechanisms^33^. It also correlates with ratings of surprise and anger. This pattern may reflect more complex responses to goal interruption or shifting, based on representations of intentions and action-outcome expectations. This accords with a previous fMRI study using verbal scenarios^41^ where appraisal features such as goal consistency, intention, or agency accounted for discriminative neural signatures in theory-of-mind networks, better than valence or arousal dimensions. More generally, these data further support appraisal theories according to which goal conduciveness and goal obstruction are key elements of emotion processing^22^.

FCP-5 is remarkable as it involves appraisal of goals (for someone else) with a striking mixture of high intensity feelings, no clear valence polarity, and prominent physiology features, particularly lump in the throat and warmth. This constellation is reminiscent of the experience of “being moved”, an emotional response whose nature and even existence is debated in the psychology literature^42^. Recent conceptual analysis in philosophy^43^, however, argued that the state of “being moved” may hold all necessary criteria to be considered a “basic” emotion, similar to traditional kinds such as fear, anger, or happiness^9^. Moreover, FCP-5 correlates with two discrete emotions of opposite valence, love and sadness, in agreement with situations typically associated with reports of “being moved” (e.g., attending a concert given by one’s child or goodbye to a soldier leaving for war). To our knowledge, this emotion has never been studied in neuroscience. Here we find not only that elements of this experience emerged as a specific FCP, but also, they distinctively mobilized midline brain areas overlapping with the DMN. Intriguingly, DMN activity is associated with introspective processing as well as access to self-related, affectively-relevant information in memory^44^ converging with the claim that “being moved” may reflect the activation of core self values^43^ and our interpretation that FCP-5 may encode social concern or “caring for others”^45^.

Finally, FCP-6 has marginal significance, but shows a distinct pattern with appraisal features related to anti-social information and motivational features of active approach, attack, and attention, positively correlated with discrete ratings of fear and disgust. A bipolar activation map with increase in VTA, VMPFC, and sensorimotor areas, but decrease in anterior insula and rostral ACC, could putatively fit well with opponent processes of approach/curiosity vs avoidance/aversion, another common dimension in classic emotion theories^6^.

Notably, the reliability of these emerging factors was verified via a bootstrap approach in which different subsets of 14 subjects out of 20 were randomly selected at every iteration of re-sampling, allowing us to test for robustness of the factors across different groups of subjects. These bootstrap results demonstrated a very small variation for each factor that implies a good relibility of factors and strong generalisability across subjects.

While our study provides several novel insights concerning the functional componential organization of emotions and their neural underpinnings, built on well-established theorizing^22^, it is not without limitations. As our approach to define latent variables (in both feature and brain space) is purely data-driven, our interpretation of FCPs necessarily implies post-hoc speculations based on extant knowledge and previous studies. This is not without a risk of reverse inferences^46^, although here brain activation patterns were confronted with predefined behavioral features and concepts derived from precise theoretical hypothesis in psychology^28^. Moreover, our findings appear consistent with other data-driven meta-analysis^17^, but go beyond usual MVPA-based approaches with pre-defined categories or dimensions that do not account for the rich variety but also similarity between emotions.

Another limitation is that our six FCPs were derived from a limited dataset (containing 40 movies) that may not embrace the whole emotion spectrum. However, to the best of our knowledge, our stimulus set is still much larger than any other standard neuroimaging studies in this field. Further, we cover a wide range of features previously validated across a variety of emotional responses^28^, and surmise that similar FCPs would occur in other emotions not elicited here (perhaps combined with additional networks).

In addition, emotions elicited by movies might most often be vicarious, in that events are not directly happening to the viewer in the real world, which may limit the generalization of our results to first-person emotions and highlights the need for using more immersive paradigms in future research to elicit first-person emotions (e.g., using games or virtual reality). However, such limitations appear even more severe in past research using pictorial stimuli such as faces or scenes. Naturalistic elicitation of emotions also implied participants were not self-monitoring and it is therefore hard to measure their level of attendance to each video clip. However, during debriefing, all participants admitted that they found all clips engaging and online monitoring through eyetracker recordings ensured they stayed awake with eyes open and lookeing at the screen. Finally, we used a single HRF function for every emotional event in movie clips to model the BOLD signal, which is a standard practice for such studies but comes with potential limitations such as neglecting carry-over, habituation, or sensitization effects. However, this should apply equally to all emotion features and would limit our analysis sensitivity rather than create spurious results.

Lastly, a striking aspect of our results is the absence of a FCP linking behavioral features and brain networks corresponding to the construct of arousal, despite this being the second major ingredient of “core affect” in classic theories of emotions^35^. This might stem from insufficient or insensitive features related to arousal and limited measures of physiological responses. Alternatively, this may support the notion that arousal does not constitute a single well-defined functional process, but encompasses variable aspects of affect, vigilance, and autonomous nervous functions^21^.

In sum, our study offers a new approach to study emotions using both theory-based and empirically validated parameters without pre-defined categories or dimensions. Our results provide new insights into the functional organization of affective processing and their relation to particular brain substrates, adding support to componential models and shedding light on neglected emotions such as “being moved”. In doing so, our work goes some way toward elucidating the constitutive ingredients of emotions and linking them to network accounts of brain function, using data-driven methodology. Nevertheless, we acknowledge that further studies are necessary to verify these findings and confirm the replicability in different contexts.

## Methods

This research has been approved by Geneva Research Ethics Committee and done according to the committee guidelines. Informed written consent was obtained from all participants.

### Emotion Elicitation

To elicit emotions in a naturalistic and dynamic manner and track component processes, rather than presenting a sequence of unrelated stimuli assigned to pre-defined categories, we used a series of 40 movie clips that were selected and validated to cover a range of different emotions, similar to those classically investigated in psychology and neuroscience. Movies provide ecologically valid stimuli as they allow for a continuous measure of emotional responses, whose nature or intensity can be influenced by context, temporal history, or expectation (beyond just the current visual or auditory inputs). The efficacy and validity of movies has been well-established in psychological studies of emotion elicitation due to these desirable characteristics^47–52^, but this approach remains scarce in neuroimaging research and limited to measures of basic dimensions (valence and arousal)^53,54^ or discrete categories of emotion (fear, sadness, etc.)^11,55^.

### Stimuli selection

To select emotionally engaging movie excerpts, we borrowed a set of 139 videos from previous researches^47,49,51,52^. All excerpts were collected in both English and French languages and, matched for duration and visual quality (the original video excerpts were in English, but our experiment was in French, so we collected the dubbed version of original clips as well). We then chose video excerpts eliciting various emotions and covering different component dimensions of interest.

To this aim, we first conducted a preliminary behavioral study where emotion assessments were made in terms of discrete emotion categories, as well as according to a set of component descriptors. Discrete emotions were rated using a modified version of the Differential Emotion Scale (14 labels: fear, anxiety, anger, shame, warm-hearted, joy, sadness, satisfaction, surprise, love, guilt, disgust, contempt, calm^56,57^), whereas the componential descriptors were assessed using a questionnaire of 39 features taken from the CoreGRID instrument^28^ (see supplementary methods, section A). The CoreGRID instrument includes 63 semantic concepts representing activity in the five major components postulated by emotion theories (appraisal, motivation, expression, physiology, feeling; see Figure 1). Our selection concerned features appropriate to emotions experienced during movie watching where events happened to characters rather than directly to the viewer in the real-world. Based on the intensity and discreteness of categorical emotion ratings, we retained 40 videos equally covering 10 different discrete emotion categories (each predominating in 4 clips, average duration 111 sec, and standard deviation of 44 sec). By using several emotions (beyond the 6 basic categories), we were able to assess a more complete range of components.

The pilot ratings and validation of video clips were obtained through a web interface using CrowdFlower, a crowdsourcing platform which allows accessing an online workforce to perform a task. The selected workforce was limited to native English speakers from USA or UK and the reward was set for an effective hourly wage of 6$. The quality control of the assessments was ensured by means of ad-hoc test questions about the content of the clip. Participants (n= 638, 358 males, mean age = 34, SD= 11) watched the full clip (on average 2.8 clips per participant), and then rated each question on how it described their feeling or experience on a 5-points Likert scale (with 1 associated to “not at all” and 5 associated to “strongly”).

### Participants in the fMRI study

Twenty right-handed volunteers (9 females, mean age 20.95, range 19 to 25 and standard deviation of 1.79) with no history of neurological or psychiatric disorders took part in the study. They were recruited via fliers and all native French speakers. All participants gave written consent according to the Geneva Research Ethics Committee guidelines. Demographic information (including age, sex, nationality, handedness, education and language speaking) and Big-Five Personality Traits (using BFI-10 questionnaire^58^) were collected prior to the experiment. There were four scanning sessions in total. Participants received a monetary reward of 40 CHF per session and a final bonus of 90 CHF if they completed all sessions (equivalent of 20 CHF per hour). For technical reasons the first session of the first participant had to be discarded from the data, and another participant only completed two sessions out of four. However, the remaining data from these two participants was used in the analysis as they included observations covering all emotion conditions.

### Experiment procedure

Participants underwent 4 testing sessions in 4 different days to complete the whole experiment. Each session started with an fMRI experiment followed by behavioral assessment. At the beginning of each fMRI session, participants completed a Brief Mood Introspection Scale (BMIS) questionnaire and got prepared for the scanning. During the fMRI scanning, they watched 10 video excerpts, belonging to 10 different pre-labelled emotion categories (in random order), ensuring to probe for component processes as equally as possible across sessions. Each video was presented once only, followed by a 30-sec washout clip (composed of geometric shapes moving over a fixed background, matched for average luminance and colour content of the preceding clip, accompanied by neutral tones sampled from the video sound track). Participants were instructed to freely feel emotions and fully appraise the affective meaning of scenes, rather than control their feeling and thoughts because of the experiment environment (see supplementary methods, section C for details).

Following each fMRI session, participants watched the same videos (whole clips) again and rated feelings and thoughts evoked by the pre-selected events in each excerpt by recalling on how they felt when they first saw it inside MRI scanner. Participants were explicitly advised to reflect on their own feelings and thoughts and not what is intended to be felt in general by watching the same event. Ratings were obtained for pre-selected events with perceptually and/or emotionally salient content (between 1 to 4 events per each video, with mean of 2.9 and duration of 12-sec per event). This allowed us to ensure ratings corresponded to a precise event, and not more global judgments about the video. Answers were given immediately after watching the emotional event by pausing the video. Emotional events were selected from a separate pilot experiment by other subjects (n = 5) who made continuous evaluations with CARMA (software for Continuous Affect Rating and Media Annotation)^59^ allowing second by second ratings (see supplementary methods, section B).

For post-fMRI ratings, participants had to choose one or two of the emotion labels (primary and secondary most felt) from the list of 10 discrete emotion categories (fear, anxiety, anger, joy, sadness, satisfied, surprise, love, disgust and calm), and to rate each of 32 CoreGRID features selected in our pilot study using a 7-point Likert scale (1 corresponding to “not at all” and 7 corresponding to “strongly”). Seven other features from the CoreGRID list concerned some of the physiological and expression measures were not included in the current analyses. Each fMRI session and behavioral session lasted for about 1h and 2h respectively (about 3h×4sessions=12h of experiment for each participant), including preparation time. Behavioral rating sessions were performed on a separate PC in a quiet room. In total, 2276 video events were rated along the 32 GRID dimensions described above (see supplementary methods, section D for details).

### Data acquisition

MRI was performed on a 3T Siemens TIM Trio scanner at the Brain & Behavior Laboratory (BBL) of the University of Geneva, with a 32-channel head coil using gradient-echo T2*-weighted echo-planar image (EPI) sequence for functional images (TR = 2000 ms, TE = 30 ms, Flip angle = 85°, FOV = 192 mm, resolution = 4 × 64, 35 axial slices, voxel size 3mm × 3mm × 3mm). High-resolution T1-weighted structural images and susceptibility weighted images (SWI) were also collected. Each video was presented during a separate MRI acquisition run to ensure independence between different stimuli. The acquisition time for each run was about 164 sec on average. Stimuli presentation and rating were controlled using Psychtoolbox-3, an interface between Matlab and computer hardware. During the fMRI session, participants watched the stimuli on an LCD screen through a mirror mounted on the head coil. The audio stream was transmitted through MRI-compatible Sensimetrics Insert earphones (model S14). Peripheral physiological measures were also collected during the fMRI session including heart activity, respiration, and electro dermal activity (EDA) using BIOPAC system. However, due to technical reasons, the EDA was missing for some of the subjects and some of the sessions. We therefore had to exclude it from further analysis. Similarly, to detect facial expressions such as smiles and frowns, we recorded electromyogram (EMG) during the full scanning sessions, but due to electromagnetic interference from other devices and poor signal overall, facial motor activity could not be reliably retrieved and was not analysed either.

### Pre-processing of fMRI data

Initial processing of the fMRI data was performed using Statistical Parametric Mapping 12 (SPM12) software (Wellcome Department of Cognitive Neurology, London, UK). The data were corrected for slice timing and motion, co-registered to high resolution structural images, and normalized to Montreal Neurologic Institute (MNI) space. Spatial smoothing was applied at 6mm and temporal data was high-pass filtered at 0.004 cut-off point. Changes in neural activation were modelled across the whole brain using the general linear model (GLM) as implemented in SPM12. For each run, the BOLD signal was modelled using multiple regressors, one per each emotional event plus one representing the washout period, which were convolved with the hemodynamic response function (HRF). The onset of each HRF is aligned with the beginning of each emotional events. Six motion parameters (translations in *x*, *y* and *z*, roll, pitch and yaw) were also added to account for nuisance effects. No mask was applied to the GLM estimations and data from whole brain was used in the later analysis. For each emotional event, a contrast map between the emotional event and washout βs was computed and then used as the differential neural marker of the corresponding emotional experience(s) associated with this event.

### Physiology signal pre-processing

All physiology signals (heart pulsation, respiratory and EDA) were acquired throughout the scanning sessions at 5000Hz sampling rate. To pre-process this data, we used AcqKnowledge 4.2 and MATLAB. Signal artefacts and signal losses were corrected manually using Endpoint function in AcqKnowledge software, which interpolates the values between two selected points. Then signals were downsampled to 120Hz and a comb-pass filter was applied to remove the scanner artifacts. The heart activity signal was filtered with a band-pass filter between 1Hz-40Hz and the heart rate (HR) was computed using peak detection technique and was converted to beats per minute. Results were controlled to make sure that the estimated HR is in the normal range of 60-100 beats per minute and otherwise corrected manually. All automatically detected peaks were verified visually and any misdetection was corrected manually. The respiration signal was band-pass filtered between 0.05Hz-1Hz and similar to heart signal, respiration rate (RR) was estimated. The EDA signal was also processed, but finally discarded from analyses due to a large portion of missing values. Average HR and RR for emotional events were computed and then treated similarly as the rest of the GRID descriptors to assess neural effects associated with increases or decreases of HR and RR.

### Component Process Model and CoreGRID

We based our analysis of behavioral and fMRI data on a previously established model^27^ assuming four different functional components, including (1) a motivational component that causes changes in action tendencies, (2) an expression component that implicates changes in motor behavior and action, (3) a physiological component that corresponds to changes in peripheral autonomic activity, and finally (4) a feeling component that reflects the conscious experience concomitant to changes in all other components^60^. A series of 144 descriptors for distinct features of each of these components has been defined in the GRID instrument^28^ in order to cover various emotional experiences according to their frequent use in the literature, cross-cultural adaptability, and common occurrence in self-reports. In our study, we used the CoreGRID instrument, a validated brief version of the GRID encompassing 63 semantic items^28^, among which we selected those applying to our experimental paradigm and compatible with a third-person perspective of emotional events. These features and their links with main components are listed in Table 1.

### Hierarchical clustering

To analyse similarity/dissimilarity between discrete emotions in terms of their profile of component features, we applied a hierarchical clustering analysis. To define this componential profile, we computed the average value of CoreGRID items for each discrete emotion class (emotion class of each rating was based on its primary emotion category) which represents the class centroids. Hierarchical clustering allows for grouping similar items into one cluster and merges pairs of clusters as it moves up the hierarchy. One advantage of such algorithm is the possibility to interpret the similarity at different levels. Here, we used squared Euclidian distance as the similarity measure between clusters (Ward’s method).

### Exploratory Factor Analysis

We applied factor analysis to all GRID items from all 4 components to find the underlying commonality across different items, allowing us to compare our dataset with previous work using similar analyses^29^. Based on Kaiser’s criterion, 6 factors were selected (see section “Underlying Factors”, which accounted for 48% of total variance, and applied orthogonal varimax rotation to simplify the expression of a particular factor in terms of just few major items. The interpretation of each factor is based on its relationship with specific GRID item-set.

### Partial Least Square Correlation

To identify consistent patterns of covariations among component features and concomitant changes in brain activity patterns, we employed Partial Least Square Correlation (PLSC), a multivariate statistical modelling technique that extracts the commonalities between neural activity and behavior through an intermediate representation of latent variables^30^. In this method, response and independent variables are projected to a new space of latent variables, such that the latter has the maximal covariance. Here, we included the behavioral ratings on all 32 CorGRID items plus 2 physiology measures (HR and RR) on one hand, and fMRI data from whole brain obtained for all emotional events across all movies (n=2276) on the other hand. In this analysis, BOLD activity from *V* voxels are stored in matrix *X_M×V_* with *M* rows as the number of observations, and ratings of emotion experience are stored in matrix *Y_M×N_* where *N* is the number of behavior descriptors. The relation between *X* and *Y* is stored in a correlation matrix *R* as *R* = *Y^T^X*, which is then submitted to Singular Value Decomposition (SVD) to obtain three different matrices as *R* = *U*Δ*V^T^*, where *U* and *V* represent saliences values (loadings) for *Y* and *X* respectively. Latent variables are computed as *L_X_* = *XV* and *L_Y_* = *YU* that model the relationship between BOLD signal and behavioral data^61^. We assessed the significance of latent variables with permutation tests^62^ (1000 iterations) and latent variables with p-value < 0.01 were retained for interpretation.

We also verified our PLSC results in terms of statistical significance and reliability using independent methodology based on permutation tests and bootstrapping, respectively. Because standard cross-validation techniques or power analysis are not applicable to the PLSC method, the optimal sample size and generalizability of results cannot be calculated solely based on the number of subjects or model parameters, but should instead be tested based on the precision of the model estimation and its standard error^63^. A robust method to estimate the precision of estimates is through resampling methods like Monte Carlo simulation, with the most popular approach based on bootstrap sampling ^62^. In our study, we took a conservative bootstrap approach, in which we limited the number of subjects used at every iteration of bootstrap to be ^~^70% of participants. This guarantees that data from ^~^30% of the subjects are excluded at every iteration and the estimates are solely based on the resampled data from a portion of subjects. As can be inferred, this method also implicitly examines the generalisability of the method to different set of samples and allowed us to estimate the robustness of our results. To do so, for each bootstrap sampling iteration, the PLSC procedure was repeated (on a set of 14 randomly selected subjects (about 70% of the subjects)) and the variance of each element of each latent variable was computed over 1000 iterations. The stability scores *S_u_i__*. and *S_v_i__*. for the *i*th elements of factors *u* and *v* are obtained as 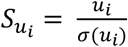 and 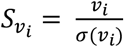, where *σ*(*u_i_*) and *σ*(*v_i_*) denote standard errors for *u_i_* and *v_i_* where *σ*(*u_i_*) and *σ*(*v_i_*) denote standard errors for *u_i_* and *v_i_* respectively. Stability scores higher than 2.5 or lower than −2.5 (corresponding to p<0.01) were considered as significant, indicating voxels that reliably respond to a particular condition. Because resampling methods can cause axis rotation and alter the order of latent variables, we used Procrustes rotation^64^ to correct for such effect. This approach allowed us to identify voxels with highest loadings for each latent variable, and emotion features contributing most to each of these latent variables. Results thus delineate distinct brain-wide networks and the component feature profiles associated with their activation, presumably underlying different dimensions of emotion over various episodes. Importantly, this patterning is obtained in a purely data-driven manner, without assigning features to particular components or appraisals, and without grouping them according to pre-defined discrete emotion categories. Please also note that our participant sample size is similar to other studies exploring social appraisal features with a smaller stimulus dataset^41^ and much larger than in other recent work using machine learning approaches with a similar comprehensive video dataset (e.g., n= 5 participant in Hirokawa et al.^65^).

## Acknowledgements

This research was supported by a grant from the Swiss National Science Foundation (SNF Sinergia No. 180319) and the National Centre of Competence in Research (NCCR) Affective Sciences (under grant No. 51NF40-104897) and it was conducted on the imaging platform at the Brain and Behavior Lab (BBL) and benefited from the support of the BBL technical staff. The authors also wish to acknowledge the great help of Maëlan Menétrey in data collection and extend their gratitude to Klaus Scherer for his insightful comments and inspiration.

## Supplementary information

### Supplementary Methods

#### A. CoreGRID descriptor selection

The CoreGRID instrument includes 63 semantic concepts capturing descriptors relevant to the five major component in the component model^1^. Several concepts in this instrument postulate a first-person perspective (where the participant is the subject of an action or is actively involved in a real situation with potential implications to oneself), and were therefore not applicable to our study where the participant is observing events in a movie. Some example of such descriptors is as follows:

- “… important and relevant for my goals”
- “… caused by my own behaviour”
- “… I spoke slower/faster”
- “… I wanted to handover the initiatives to someone else”
- “… I had resources to avoid or modify consequences”

Hence all such descriptors were eliminated from the list, which resulted in 39 items to be used for ratings (see Supplementary Table 1 for the list of descriptors in the preliminary stimuli selection study). In addition, the item “speech disturbances” was also eliminated for the fMRI study as participants are restricted in moving or talking inside the scanner. Moreover, for 4 physiology descriptors including “heartbeat getting faster”, “breathing slowing down”, “felt warm”, and “sweat”, and 2 facial expressions of “smiling” and “frowning”, we decided to use objective measures through sensors, and therefore these 6 descriptors were included in the preliminary selection study but excluded from subjective ratings in the fMRI study. Thus, in the final experiment, participants had to judge their affective experience in terms of the remaining 32 descriptors from the GRID.

**Supplementary Table 1:** Selected GRID descriptors. This table shows the list of GRID descriptors selected for the preliminary stimulus selection study. For the final fMRI study, 38 items were retained out of which 6 were collected objectively through sensors (but due to technical reasons, only HR and RR could be used in final analysis, see main text).

|  | **GRID selected descriptors** | *Component* |
| --- | --- | --- |
|  | While watching this movie, did you... |  |
| 1 | think it was incongruent with your standards/ideas? | Appraisal |
| 2 | feel that the event was unpredictable? | Appraisal |
| 3 | feel the event occurred suddenly? | Appraisal |
| 4 | think the event was caused by chance? | Appraisal |
| 5 | think that the consequence was predictable? | Appraisal |
| 6 | feel it was unpleasant for someone else? | Appraisal |
| 7 | think it was important for somebody's goal or need? | Appraisal |
| 8 | think it violated laws/social norms? | Appraisal |
| 9 | feel in itself was unpleasant for you? | Motivation |
| 10 | want the situation to continue? | Motivation |
| 11 | feel the urge to stop what was happening? | Motivation |
| 12 | want to undo what was happening? | Motivation |
| 13 | lack the motivated to pay attention to the scene? | Motivation |
| 14 | want to destroy s.th.? | Motivation |
| 15 | want to damage, hit or say s.th. that hurts? | Motivation |
| 16 | want to tackle the situation and do s.th.? | Motivation |
| 17 | have a feeling of lump in the throat? | Physiology |
| 18 | have stomach trouble? | Physiology |
| 19 | experience muscles tensing? | Physiology |
| 20 | feel warm? | Physiology |
| 21 | sweat? | Physiology |
| 22 | feel heartbeat getting faster? | Physiology |
| 23 | feel breathing getting faster? | Physiology |
| 24 | feel breathing slowing down? | Physiology |
| 25 | produce abrupt body movement? | Expression |
| 26 | close your eyes? | Expression |
| 27 | press lips together? | Expression |
| 28 | have the jaw drop? | Expression |
| 29 | show tears? | Expression |
| 30 | have eyebrow go up? | Expression |
| 31 | smile? | Expression |
| 32 | frown? | Expression |
| 33 | produce speech disturbances? | Feeling |
| 34 | feel good? | Feeling |
| 35 | feel bad? | Feeling |
| 36 | feel calm? | Feeling |
| 37 | feel strong? | Feeling |
| 38 | feel an intense emotional state? | Feeling |
| 39 | experience an emotional state for a long time? | Feeling |

**Supplementary Table 2:** High saliency brain areas. This table lists the name of the films for each clip, its duration, the original emotion label as collected in another study and number of events selected in each clip.

| Films | Duration (s) | Related Emotion | Segments |
| --- | --- | --- | --- |
| 12 Years A Slave | 120 | Anger | 4 |
| 28 Days Later | 143 | Fear | 4 |
| 28 Days Later | 49 | Anxiety | 4 |
| 28 Days Later | 125 | Fear | 4 |
| A Fish Called Wanda | 173 | Joy | 4 |
| American History X | 77 | Anxiety | 4 |
| Bambi | 86 | Love | 2 |
| Batman Returns | 73 | Surprise | 3 |
| Cry Freedom | 132 | Disgust | 4 |
| Dangerous Minds | 128 | Sadness | 2 |
| Edward Scissorhands | 93 | Calm | 4 |
| Forrest Gump | 121 | Love | 3 |
| Ghost | 215 | Love | 4 |
| Hotel Rwanda | 209 | Sadness | 4 |
| Hotel Rwanda | 96 | Sadness | 2 |
| Hotel Rwanda | 97 | Anger | 3 |
| Kill Bill 1 | 68 | Surprise | 2 |
| Kill Bill 1 | 144 | Surprise | 2 |
| Life Is Beautiful | 127 | Anger | 3 |
| Life Is Beautiful | 105 | Love | 2 |
| Love Actually | 76 | Joy | 3 |
| Love Actually | 104 | Joy | 4 |
| Love Actually | 28 | Satisfaction | 1 |
| Mr. Bean's Holiday | 128 | Joy | 4 |
| My Girl | 146 | Sadness | 2 |
| Planet Earth | 66 | Calm | 3 |
| Pride And Prejudice | 91 | Calm | 3 |
| Remember The Titans | 132 | Satisfaction | 3 |
| Schindler's List | 115 | Disgust | 2 |
| Scream 2 | 215 | Fear | 4 |
| Terminator 2 | 131 | Satisfaction | 4 |
| The Dentist | 56 | Fear | 2 |
| The Fly | 81 | Anxiety | 3 |
| The Pianist | 81 | Anger | 2 |
| The Pianist | 117 | Anxiety | 2 |
| There Is Something About Mary | 175 | Surprise | 4 |
| The Shining | 85 | Satisfaction | 2 |
| Trainspotting | 104 | Disgust | 3 |
| Trainspotting | 62 | Disgust | 1 |
| Wild Alaska | 62 | Calm | 2 |

#### B. Emotional segment selection

Although the initial assessment to select video clips was done on the basis of global judgments, to avoid any confound of multiple events related to different appraisal components within a clip, we analysed each single emotional event separately for the final fMRI study. The selection of emotional events was semi-manual and done through a separate experiment. In that experiment, five participants were asked to watch the video clips and continuously rate the emotional content using the CARMA tool (software for Continuous Affect Rating and Media Annotation)^2^. The ratings were quantified on a range from 0 to 100 and, for each participant, ratings above the mean value 32 (the value has been rounded) were selected as emotional moments. Events that were annotated as emotional by a majority of participants (3 or more participants) were selected as emotional segments. The start and end of each segment was manually adjusted to have the same length across all participants in the final fRMI study. Supplementary Table 2 lists the name of the films for each clip and the number of emotional events selected in that clip.

This selection of emotional events resulted in a non-uniform distribution of discrete emotions, as depicted in Figure 3a. This is unlike the global judgments made on the initial video clip dataset and used to select the final experimental movie list, indicating that such global assessment of video clips (as done in may studies) may not necessarily apply to all its single events within the clip, and thus highlighting the importance of focusing on short segments to evaluate felt emotions. Please note that this non-uniform distribution might be considered as one limitation of the current study but does not necessarily affect the main results since our main goal is to cover a wide range of componential space, not to elicit or compare specific discrete emotions.

#### C. fMRI session

Each fMRI session started off by giving instructions regarding the experiment and fMRI acquisition protocol. Next, participants had to fill in the required forms and completed a 16-item Brief Mood Introspection Scale (BMIS) mood questionnaire^3^. When the participant was ready, (s)he entered the scanner and the physiology collecting devices, EMG, headphones, and eye-tracker were set up. After checking all physiology signals and calibrating the eye-tracker, the actual acquisition started. Each video clip was played during one single run and each run lasted ^~^164s including a short initial preparation time and final washout clip. There was an interval of ^~^30 s between consecutive runs. Supplementary Figure 1 illustrates the sequence of events inside the fMRI scanner. On average, the overall time inside the scanner for each session was about 32 minutes, excluding the setup time, calibration and structural MRI acquisition.

**Supplementary Figure 1:**
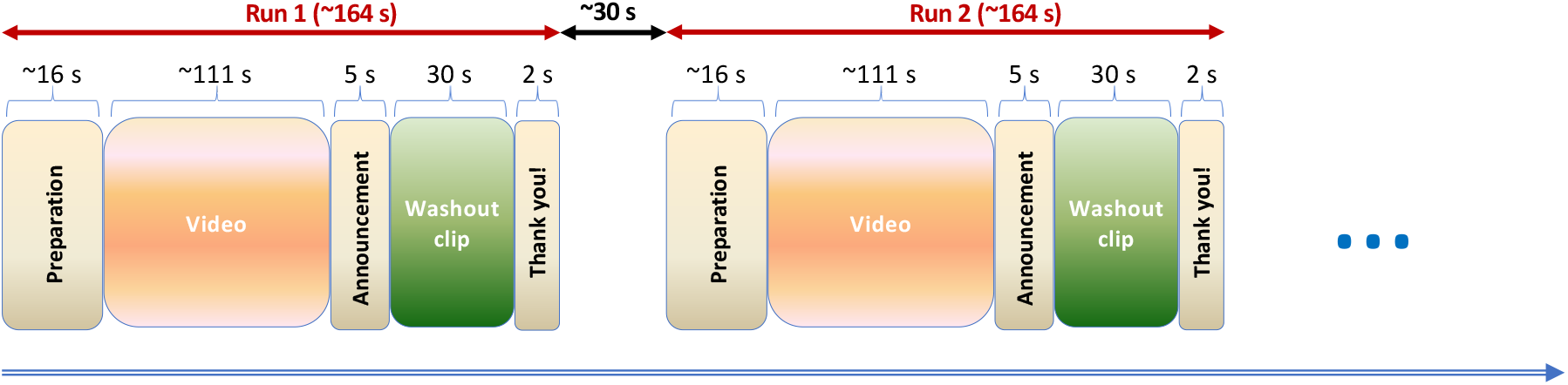
fMRI experiment design. This figure shows the design of experiment when participants where inside the scanner and the average time course of every phase. There was about 30 seconds intervals between two consecutive runs.

#### D. Behavioral session

Similar to fMRI session, at the beginning of each behavioral session, participants were briefed on the procedure to evaluate their felt emotions. The experiment started by asking them to answer a 10-item Big Five Inventory (BFI) personality questionnaire^4^, followed by playing the same video clips as those seen in the fMRI session. Emotional segments were highlighted by a red frame to notify the participant of the video segment/event, (s) he hadto **a**ssess. After each emotional segment, the video was paused, and the participant had to answer 32 questions covering the GRID component items and 2 questions regarding the two most dominant discrete emotions they felt. Each behavioral session took on average about 110 minutes excluding the instruction time. Supplementary Figure 2 illustrates the behavioral rating experiment.

**Supplementary Figure 2:**
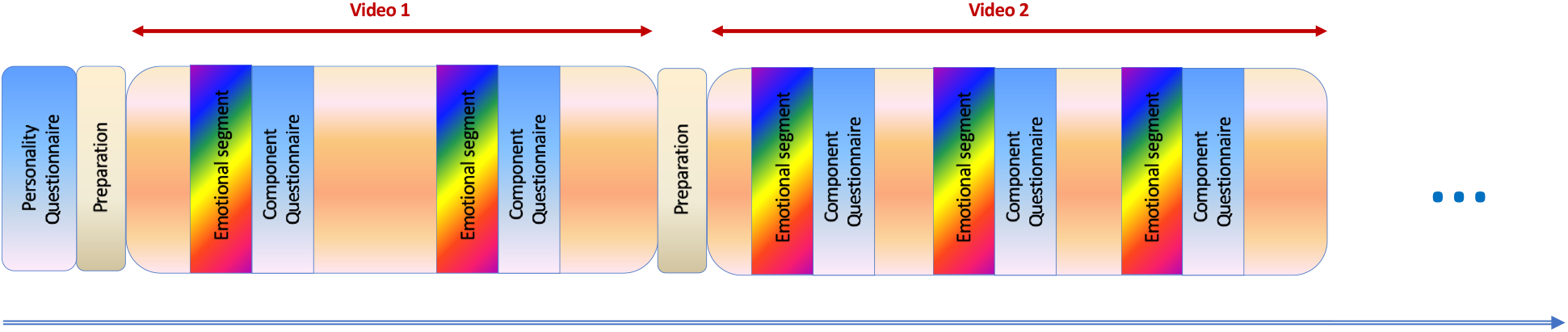
Behavioral experiment design. This figure shows the design of experiment when participants were outside fMRI scanner and assessed their subjective emotion experience for every highlighted emotional segment in movies.

### Supplementary Results

#### A. Brain Saliencies

The partial least square correlation method estimates saliency values for each voxel in the brain. To summarise the list of regions with high saliencies we have used Automated Anatomical Labelling (AAL) atlas^5^ and reported the results in Supplementary Table 3. The reported values are the average saliencies across the voxels of a given area (with cut-off point 3 for positive values and −3 for negative values) and therefore may artificially reduce or augment the apparent importance of some regions relative to others. Regions that do not show any high saliencies for any of the latent variables are not listed

**Supplementary Table 3:** High saliency brain areas. This table lists the brain areas (excluding brain stem) as defined in AAL atlas with high average saliencies (>3 or <-3) corresponding to each of the 6 latent variables. Average values across voxels of a given region may artificially increase or decrease the apparent importance of some regions relative to others.

| Region | LV1 | LV2 | LV3 | LV4 | LV5 | LV6 |
| --- | --- | --- | --- | --- | --- | --- |
| 'Precentral_L' | - | 3.52 | 3.27 | - | - | - |
| 'Precentral_R' | - | - | 3.46 | -3.38 | - | - |
| 'Frontal_Sup_L' | - | - | - | - | - | 3.14 |
| 'Frontal_Sup_R' | - | - | - | - | - | 3.32 |
| 'Frontal_Sup_Orb_L' | - | - | - | -3.37 | - | - |
| 'Frontal_Sup_Orb_R' | 3.25 | 3.34 | -3.24 | - | - | - |
| 'Frontal_Mid_L' | - | 3.17 | - | -3.11 | 3.42 | - |
| 'Frontal_Mid_R' | - | - | - | -3.28 | - | - |
| 'Frontal_Mid_Orb_L' | 3.60 | - | - | - | - | - |
| 'Frontal_Mid_Orb_R' | 3.26 | - | - | -3.15 | - | - |
| 'Frontal_Inf_Oper_L' | -3.39 | 3.49 | - | - | - | -3.11 |
| 'Frontal_Inf_Oper_R' | -3.28 | - | 3.18 | - | - | -3.16 |
| 'Frontal_Inf_Tri_L' | - | 3.26 | - | - | 3.38 | - |
| 'Frontal_Inf_Tri_R' | - | 3.22 | - | - | - | - |
| 'Frontal_Inf_Orb_R' | - | - | - | - | 3.10 | -3.31 |
| 'Rolandic_Oper_L' | -3.47 | - | 3.77 | - | - | - |
| 'Rolandic_Oper_R' | -3.25 | - | 4.10 | - | - | - |
| 'Supp_Motor_Area_L' | - | 3.16 | 3.28 | - | - | - |
| 'Supp_Motor_Area_R' | - | 3.14 | 3.17 | -3.55 | - | - |
| 'Olfactory_L' | - | - | - | - | 3.46 | - |
| 'Olfactory_R' | - | - | - | - | 3.44 | - |
| 'Frontal_Sup_Medial_L' | - | 3.42 | - | - | 3.16 | - |
| 'Frontal_Sup_Medial_R' | - | 3.48 | - | - | - | - |
| 'Frontal_Med_Orb_L' | - | - | - | -3.47 | 3.26 | 3.27 |
| 'Frontal_Med_Orb_R' | - | 3.22 | -3.14 | -3.36 | 3.57 | - |
| 'Rectus_L' | 3.74 | - | - | -3.49 | - | - |
| 'Rectus_R' | 3.49 | 3.43 | -3.33 | -3.43 | 3.34 | - |
| 'Insula_L' | -3.88 | - | 3.53 | - | 3.39 | -3.81 |
| 'Insula_R' | -3.79 | - | 3.58 | - | - | -3.64 |
| 'Cingulum_Ant_L' | - | - | - | -3.31 | 3.72 | - |
| 'Cingulum_Ant_R' | -3.16 | - | - | -3.20 | 3.69 | - |
| 'Cingulum_Mid_L' | -3.35 | - | 3.45 | -3.14 | 3.20 | - |
| 'Cingulum_Mid_R' | -3.37 | - | - | -3.32 | 3.32 | - |
| 'Cingulum_Post_L' | - | - | - | - | 3.28 | 3.67 |
| 'Cingulum_Post_R' | -3.21 | - | -4.23 | - | 3.55 | - |
| 'Hippocampus_L' | - | - | - | - | - | 3.42 |
| 'Hippocampus_R' | -3.57 | - | - | - | - | 3.26 |
| 'ParaHippocampal_L' | - | - | - | - | 3.15 | - |
| 'Amygdala_L' | -3.19 | - | - | - | - | - |
| 'Amygdala_R' | - | - | - | - | - | 3.36 |
| 'Calcarine_L' | -3.24 | 3.11 | - | - | - | - |
| 'Cuneus_L' | -3.14 | - | - | - | - | - |
| 'Cuneus_R' | -3.61 | - | - | - | - | - |
| 'Lingual_L' | -3.43 | - | - | - | - | - |
| 'Lingual_R' | -3.22 | - | - | - | - | - |
| 'Occipital_Sup_L' | -3.40 | - | - | -3.36 | - | - |
| 'Occipital_Sup_R' | - | - | - | -3.73 | -3.18 | - |
| 'Occipital_Mid_L' | -3.96 | - | - | - | - | - |
| 'Occipital_Mid_R' | -3.80 | - | - | - | -3.18 | - |
| 'Occipital_Inf_L' | -3.55 | - | - | - | - | - |
| 'Occipital_Inf_R' | - | - | -3.16 | - | - | - |
| 'Fusiform_L' | -3.34 | - | - | - | - | - |
| 'Fusiform_R' | -3.23 | - | - | - | - | - |
| 'Postcentral_L' | - | - | 3.67 | -3.18 | - | 3.15 |
| 'Postcentral_R' | - | - | 3.69 | -3.67 | - | 3.43 |
| 'Parietal_Sup_L' | -3.29 | - | - | -3.28 | - | - |
| 'Parietal_Sup_R' | -3.44 | - | - | -3.85 | - | 3.18 |
| 'Parietal_Inf_L' | -3.43 | - | - | -3.39 | - | - |
| 'Parietal_Inf_R' | -3.35 | - | - | -4.36 | - | - |
| 'SupraMarginal_L' | -4.87 | 3.24 | 3.15 | - | 3.72 | - |
| 'SupraMarginal_R' | -4.55 | - | - | - | - | - |
| 'Angular_L' | - | - | - | - | 4.43 | - |
| 'Angular_R' | - | - | -3.38 | -3.74 | 3.21 | - |
| 'Precuneus_L' | -3.65 | 3.17 | - | - | - | 3.17 |
| 'Precuneus_R' | -3.61 | - | -3.69 | - | - | 3.17 |
| 'Paracentral_Lobule_L' | - | - | 3.42 | -3.60 | - | 3.52 |
| 'Paracentral_Lobule_R' | - | - | 3.29 | -3.52 | - | 3.12 |
| 'Caudate_L' | -3.17 | - | - | - | - | - |
| 'Caudate_R' | -3.38 | - | - | - | - | 3.32 |
| 'Putamen_L' | -3.37 | - | - | - | 3.20 | - |
| 'Putamen_R' | -3.52 | - | 3.33 | - | 3.14 | - |
| 'Pallidum_R' | -3.52 | - | - | 3.39 | 3.20 | - |
| 'Thalamus_L' | -4.08 | - | - | - | - | - |
| 'Thalamus_R' | -4.20 | - | - | - | - | - |
| 'Heschl_L' | - | - | 3.37 | - | - | - |
| 'Heschl_R' | - | 3.30 | 3.49 | - | - | - |
| 'Temporal_Sup_L' | - | 3.57 | 3.44 | - | - | - |
| 'Temporal_Sup_R' | - | 3.81 | - | 3.11 | - | - |
| 'Temporal_Pole_Sup_L' | 3.75 | 3.34 | 3.90 | - | 3.20 | -3.18 |
| 'Temporal_Pole_Sup_R' | 3.13 | 3.25 | 3.70 | - | - | - |
| 'Temporal_Mid_L' | - | 3.52 | - | - | - | - |
| 'Temporal_Mid_R' | -3.64 | 3.87 | -3.52 | - | - | - |
| 'Temporal_Pole_Mid_L' | - | - | - | - | 3.31 | - |
| 'Temporal_Pole_Mid_R' | - | - | -3.13 | - | - | - |
| 'Temporal_Inf_L' | - | 3.18 | - | - | - | - |
| 'Temporal_Inf_R' | -3.55 | - | - | -4.04 | - | - |

#### B. Discrete emotion and latent variables correlation

To examine the similarity between each latent variable and the discrete emotion profiles, a Pearson correlation analysis was performed. Supplementary Figure 3 depicts the significance of these correlations, representing the relative implication of each LV in discrete emotion categories.

Almost all discrete emotions showed high positive or negative correlations with the first latent variable LV1 (encoding appraisals of values or valence), except for sadness and surprise that showed weaker or no significant correlation. The second latent variable LV2 (attributed to novelty) showed a strong positive correlation with surprise, but negative correlation with sadness and anger, while LV3 (interpreted as hedonic impact) was positively correlated with joy, satisfaction, love and calm, but negatively correlated with anxiety and sadness. LV4 (related to goals and intentions) was positively associated with surprise and negatively with anger. Interestingly, the fifth latent variable LV5 was found to be significantly correlated with sadness followed by high non-significant correlation with love, two discrete emotions of opposite valence (but consistent with a dimension of caring for others and social concern). And finally, the last latent variable LV6 (encoding dimensions of curiosity and active approach) showed significant positive correlation with fear followed by high non-significant correlations with anxiety and disgust. Taken together, these findings highlight that each of the different LVs identified by our data driven PLSC analysis contributed to different emotions, but to variable degrees, and also that they generally held meaningful relationships with discrete categorical labels. Importantly, however, single LVs cannot be reduced to particular emotion categories or unique orthogonal dimensions such as valence or arousal.

**Supplementary Figure 3:**
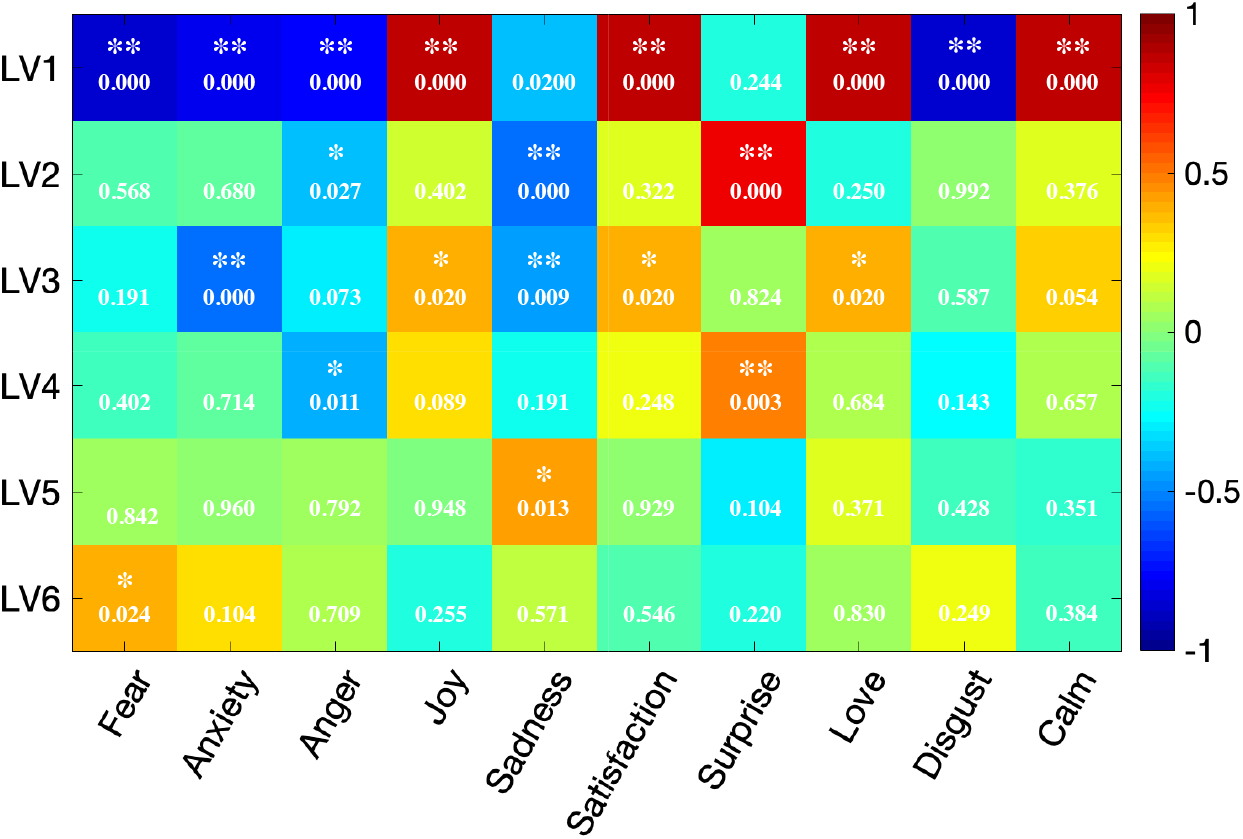
Correlation analysis. Pearson correlation coefficient between latent variables (LV) and discrete emotion categories based on individual ratings of movies (** corresponds to p-value<.01 and * corresponds to p-value<.05)

## References

1. Phelps, E. A., Ling, S. & Carrasco, M. Emotion facilitates perception and potentiates the perceptual benefits of attention. Psychol. Sci. 17, 292–299 (2006).

2. Pessoa, L. On the relationship between emotion and cognition. Nat. Rev. Neurosci. 9, 148–158 (2008).

3. Phelps, E. A. Human emotion and memory: interactions of the amygdala and hippocampal complex. Curr. Opin. Neurobiol. 14, 198–202 (2004).

4. Tambini, A., Rimmele, U., Phelps, E. A. & Davachi, L. Emotional brain states carry over and enhance future memory formation. Nat. Neurosci. 20, 271–278 (2017).

5. Damasio, A. R. Descartes’ Error: Emotion, Reason and the Human Brain. (Penguin Random House, 2006).

6. Sander, D., Grandjean, D. & Scherer, K. R. A systems approach to appraisal mechanisms in emotion. Neural Networks 18, 317–352 (2005).

7. Adolphs, R. How should neuroscience study emotions? by distinguishing emotion states, concepts, and experiences. Soc. Cogn. Affect. Neurosci. 12, 24–31 (2017).

8. Meaux, E. & Vuilleumier, P. Emotion Perception and Elicitation. in Brain Mapping: An Encyclopedic Reference 79–90 (2015).

9. Ekman, P. Basic Emotions. in Handbook of Cognition and Emotion. 301–320 (Wiley, New York, 1999).

10. Russell, J. A. Core affect and the psychological construction of emotion. Psychol. Rev. 110, 145–172 (2003).

11. Saarimaki, H. et al. Discrete Neural Signatures of Basic Emotions. Cereb. Cortex 26, 2563–2573 (2015).

12. Kragel, P. A. & LaBar, K. S. Multivariate pattern classification reveals autonomic and experiential representations of discrete emotions. Emotion 13, 681–690 (2013).

13. Kragel, P. A. & LaBar, K. S. Multivariate neural biomarkers of emotional states are categorically distinct. Soc. Cogn. Affect. Neurosci. 10, 1437–1448 (2015).

14. Kassam, K. S., Markey, A. R., Cherkassky, V. L., Loewenstein, G. & Just, M. A. Identifying Emotions on the Basis of Neural Activation. PLoS One 8, (2013).

15. Wager, T. D. et al. A Bayesian Model of Category-Specific Emotional Brain Responses. PLoS Comput. Biol. 11, 1–27 (2015).

16. Lindquist, K. A., Wager, T. D., Kober, H., Bliss-Moreau, E. & Barrett, L. F. The brain basis of emotion: A meta-analytic review. Behav. Brain Sci. 35, 121–143 (2012).

17. Kober, H. et al. Functional grouping and cortical-subcortical interactions in emotion: A meta-analysis of neuroimaging studies. Neuroimage 42, 998–1031 (2008).

18. Pool, E., Brosch, T., Delplanque, S. & Sander, D. Attentional bias for positive emotional stimuli: A meta-analytic investigation. Psychol. Bull. 142, 79–106 (2016).

19. Berridge, K. C. Affective valence in the brain: modules or modes? Nat. Rev. Neurosci. 20, 225–234 (2019).

20. Haj-Ali, H., Anderson, A. K. & Kron, A. Comparing three models of arousal in the human brain. Soc. Cogn. Affect. Neurosci. 15.1, 1–11 (2020).

21. Satpute, A. B., Kragel, P. A., Barrett, L. F., Wager, T. D. & Bianciardi, M. Deconstructing arousal into wakeful, autonomic and affective varieties. Neurosci. Lett. 693, 19–28 (2019).

22. Scherer, K. R. Emotions are emergent processes: they require a dynamic computational architecture. Philos. Trans. R. Soc. Lond. B. Biol. Sci. 364, 3459–74 (2009).

23. Scherer, K. R. & Moors, A. The Emotion Process: Event Appraisal and Component Differentiation. Annu. Rev. Psychol. 70, 719–745 (2019).

24. Ellsworth, P. C. & Scherer, K. R. Appraisal processes in emotion. Handbook of affective sciences 572–595 (2003).

25. Damasio, A. R. et al. Subcortical and cortical brain activity during the feeling of self-generated emotions. Nat. Neurosci. 3, 1049–56 (2000).

26. Nummenmaa, L. et al. Emotions promote social interaction by synchronizing brain activity across individuals. Proc. Natl. Acad. Sci. 109, 9599–9604 (2012).

27. Scherer, K. R. Appraisals considered as a process of multilevel sequential checking. in Appraisal Processes in Emotion: Theory, Methods, Research 92–120 (2001).

28. Fontaine, J., Scherer, K. & Soriano, C. Components of Emotional Meaning: A sourcebook. (Oxford University Press, 2013).

29. Fontaine, J. R. J., Scherer, K. R., Roesch, E. B. & Ellsworth, P. C. The World of Emotions Is Not Two-Dimensional. Psychol. Sci. 18, 1050–1057 (2007).

30. Krishnan, A., Williams, L. J., McIntosh, A. R. & Abdi, H. Partial Least Squares (PLS) methods for neuroimaging: A tutorial and review. Neuroimage 56, 455–475 (2011).

31. Raichle, M. E. The Brain’s Default Mode Network. Annu. Rev. Neurosci. 38, 433–447 (2015).

32. Amodio, D. M., Frith, C. D. & Frith, C. D. (2006) Meeting of minds: the medial frontal cortex and social cognition. 183–207 (2016). doi:10.4324/9781315630502-18

33. Corbetta, M. & Shulman, G. L. Control of goal-directed and stimulus-driven attention in the brain. Nat. Rev. Neurosci. 3, 201–215 (2002).

34. Harrison, N. A., Kreibig, S. D. & Critchley, H. A two way road. in The Cambridge handbook of human affective neuroscience 82–106 (2013).

35. Russell, J. A. & Barrett, L. F. Core affect, prototypical emotional episodes, and other things called emotion: Dissecting the elephant. J. Pers. Soc. Psychol. 76, 805–819 (1999).

36. Berridge, K. C. Wanting and Liking: Observations from the Neuroscience and Psychology Laboratory. Inquiry 52, 378–398 (2009).

37. Caruana, F. et al. Mirth and laughter elicited by electrical stimulation of the human anterior cingulate cortex. Cortex 71, 323–331 (2015).

38. Delplanque, S. et al. Sequential unfolding of novelty and pleasantness appraisals of odors: Evidence from facial electromyography and autonomic reactions. Emotion 9, 316–328 (2009).

39. Kron, A., Pilkiw, M., JBanaei, asmin, Goldstein, A. & Anderson, A. K. Are valence and arousal separable in emotional experience? Emotion 15, (2015).

40. Schwartz, C. E. et al. Differential amygdalar response to novel versus newly familiar neutral faces: a functional MRI probe developed for studying inhibited temperament. Biol. Psychiatry 53, 854–862 (2003).

41. Skerry, A. E. & Saxe, R. Neural Representations of Emotion Are Organized around Abstract Event Features. Curr. Biol. 25, 1945–1954 (2015).

42. Zickfeld, J. H., Schubert, T. W., Seibt, B. & Fiske, A. P. *Moving* Through the Literature: What Is the Emotion Often Denoted *Being Moved?* Emot. Rev. 11, 123–139 (2019).

43. Cova, F. & Deonna, J. A. Being moved. Philos. Stud. 169, 447–466 (2014).

44. D’Argembeau, A. & Van der Linden, M. Influence of Affective Meaning on Memory for Contextual Information. Emotion 4, 173–188 (2004).

45. Helm, B. W. Love, Friendship, and the Self: Intimacy, Identification, and the Social nature of persons. (Oxford University Press, 2010).

46. Poldrack, R. A. Can cognitive processes be inferred from neuroimaging data? Trends Cogn. Sci. 10, 59–63 (2006).

47. Gross, J. J. & Levenson, R. W. Emotion Elicitation using Films. Cogn. Emot. 9, 87–108 (1995).

48. Philippot, P. Inducing and Assessing Differentiated Emotion-Feeling States in the Laboratory. Cogn. Emot. 7, 171–193 (1993).

49. Schaefer, A., Nils, F., Philippot, P. & Sanchez, X. Assessing the effectiveness of a large database of emotion-eliciting films: A new tool for emotion researchers. Cogn. Emot. 24, 1153–1172 (2010).

50. Samson, A. C., Kreibig, S. D., Soderstrom, B., Wade, A. A. & Gross, J. J. Eliciting positive, negative and mixed emotional states: A film library for affective scientists. Cogn. Emot. 30, 827–856 (2016).

51. Gabert-Quillen, C. A., Bartolini, E. E., Abravanel, B. T. & Sanislow, C. A. Ratings for emotion film clips. Behav. Res. Methods 47, 773–787 (2015).

52. Soleymani, M., Chanel, G., Kierkels, J. J. M. & Pun, T. Affective ranking of movie scenes using physiological signals and content analysis. Proceeding 2nd ACM Work. Multimed. Semant. 32–39 (2008).

53. Sabatinelli, D. et al. Emotional perception: Meta-analyses of face and natural scene processing. Neuroimage 54, 2524–2533 (2011).

54. Lahnakoski, J. M. et al. Naturalistic fMRI Mapping Reveals Superior Temporal Sulcus as the Hub for the Distributed Brain Network for Social Perception. Front. Hum. Neurosci. 6, 233 (2012).

55. Tettamanti, M. et al. Distinct pathways of neural coupling for different basic emotions. Neuroimage 59, 1804–1817 (2012).

56. Izard, C. E., Libero, D. Z., Putnam, P. & Haynes, O. M. Stability of emotion experiences and their relations to traits of personality. J. Pers. Soc. Psychol. 64, 847–860 (1993).

57. McHugo, G. J., Smith, C. A. & Lanzetta, J. T. The structure of self-reports of emotional responses to film segments. Motiv. Emot. 6, 365–385 (1982).

58. Rammstedt, B. & John, O. P. Measuring personality in one minute or less: A 10-item short version of the Big Five Inventory in English and German. J. Res. Pers. 41, 203–212 (2007).

59. Girard, J. M. CARMA: Software for continuous affect rating and media annotation. J. Open Res. Softw. 2, (2014).

60. Scherer, K. R. The dynamic architecture of emotion: Evidence for the component process model. Cogn. Emot. 23, 1307–1351 (2009).

61. Abdi, H. & Williams, L. J. Partial Least Squares Methods: Partial Least Squares Correlation and Partial Least Square Regression. in Computational Toxicology (Humana Press, Totowa, NJ, 2012).

62. Efron, B. & Tibshirani, R. Bootstrap Methods for Standard Errors, Confidence Intervals, and Other Measures of Statistical Accuracy. Stat. Sci. 54–75 (1986).

63. Marcoulides, G. A. & Chin, W. W. You Write, but Others Read: Common Methodological Misunderstandings in PLS and Related Methods. Springer Proc. Math. Stat. 56, 31–64 (2013).

64. Ten Berge, J. M. F. Orthogonal procrustes rotation for two or more matrices. Psychometrika 42, 267–276 (1977).

65. Horikawa, T., Cowen, A. S., Keltner, D. & Kamitani, Y. The Neural Representation of Visually Evoked Emotion Is High-Dimensional, Categorical, and Distributed across Transmodal Brain Regions. iScience 101060 (2020).

## Supplementary references

1. Fontaine, J., Scherer, K. & Soriano, C. Components of Emotional Meaning: A sourcebook. (Oxford University Press, 2013).

2. Girard, J. M. CARMA: Software for continuous affect rating and media annotation. J. Open Res. Softw. 2, (2014).

3. Mayer, J. & Gaschke, Y. The Brief Mood Introspection Scale (BMIS). UNH Personal. Lab (1988).

4. Anderson, C. L. et al. Measuring personality in one minute or less: A 10-item short version of the Big Five Inventory in English and German. MIS Q. 41, 273–284 (2010).

5. Tzourio-Mazoyer, N. et al. Automated anatomical labeling of activations in SPM using a macroscopic anatomical parcellation of the MNI MRI single-subject brain. Neuroimage 15, 273–289 (2002).

